# A live-cell screen for altered Erk dynamics reveals principles of proliferative control

**DOI:** 10.1101/675736

**Authors:** Alexander G. Goglia, Maxwell Z. Wilson, Jillian Silbert, Lena P. Basta, Siddhartha G. Jena, Danelle Devenport, Jared E. Toettcher

**Affiliations:** Department of Molecular Biology, Princeton University, Princeton, NJ 08544, USA

**Keywords:** Ras, Erk, MAP kinase, cell signaling, dynamics, drug screen

## Abstract

Complex, time-varying responses have been observed widely in cell signaling, but how specific dynamics are generated or regulated is largely unknown. One major obstacle has been that high-throughput screens for identifying pathway components are typically incompatible with the live-cell assays used to monitor dynamics. Here, we address this challenge by performing a drug screen for altered Erk signaling dynamics in primary mouse keratinocytes. We screened a library of 429 kinase inhibitors, monitoring Erk activity over 5 h in more than 80,000 single live cells. The screen revealed both known and uncharacterized modulators of Erk dynamics, including inhibitors of non-EGFR receptor tyrosine kinases (RTKs) that increased Erk pulse frequency and overall activity. Using drug treatment and direct optogenetic control, we demonstrate that drug-induced changes to Erk dynamics alter the conditions under which cells proliferate. Our work opens the door to high-throughput screens using live-cell biosensors and reveals that cell proliferation integrates information from Erk dynamics as well as additional permissive cues.

## Introduction

Animal cells must respond to a large number of external cues to function appropriately during development and adult tissue homeostasis. To that end, a typical mammalian cell is endowed with hundreds of distinct receptors, yet only a few signaling pathways downstream of these receptors are tasked with responding to these many inputs. For instance, the 58 human receptor tyrosine kinases (RTKs) activate on the order of ten intracellular pathways (e.g., Ras/Erk, PI3K/Akt, Src, PLC*γ*, calcium), yet can trigger diverse downstream cellular responses in developing and adult tissues (Downward, 2001; Lemmon and Schlessinger, 2010). Cells are thus faced with the challenge of accurately transmitting information from many upstream inputs using only a few ‘wires’ or signal transduction pathways.

One resolution to this paradox comes in the form of dynamic regulation. Two receptors may trigger different time-varying responses from a single pathway, which can then be interpreted into distinct fates (Marshall, 1995). Indeed, many core mammalian signaling pathways have now been observed to generate complex, time-varying signaling behaviors in response to certain input stimuli (Purvis and Lahav, 2013). A growing body of evidence suggests that these dynamics are relevant to normal cell function: Erk and p53 pulses have been observed *in vivo* with similar timescales to those found in cultured cells (de la Cova et al., 2017; Hamstra et al., 2006; Hiratsuka et al., 2015) and disease-associated Ras/Erk pathway mutations can alter dynamics in a manner that affects cell proliferation (Bugaj et al., 2018). Yet it largely remains an open question how signaling dynamics are generated, regulated, and interpreted. We reasoned that a powerful tool to address this question would be to conduct a screen using signaling dynamics as a phenotypic readout (Behar et al., 2013; Stewart-Ornstein and Lahav, 2017).

Such a screen could potentially generate many useful insights. Perturbations to certain signaling nodes might eliminate dynamics altogether (**Figure 1A**, top), identifying essential components for producing time-varying responses. Others might switch dynamics between normal and diseased states (**Figure 1A**, middle), suggesting intervention points to correct aberrant signaling. Finally, some perturbations might result in hitherto-unobserved signaling dynamics (**Figure 1A**, bottom). These novel dynamic responses are analogous to classical mutant phenotypes, which can be highly informative despite (or because of) their differences from what is normally observed (Nusslein-Volhard and Wieschaus, 1980).

**Fig. 1.**
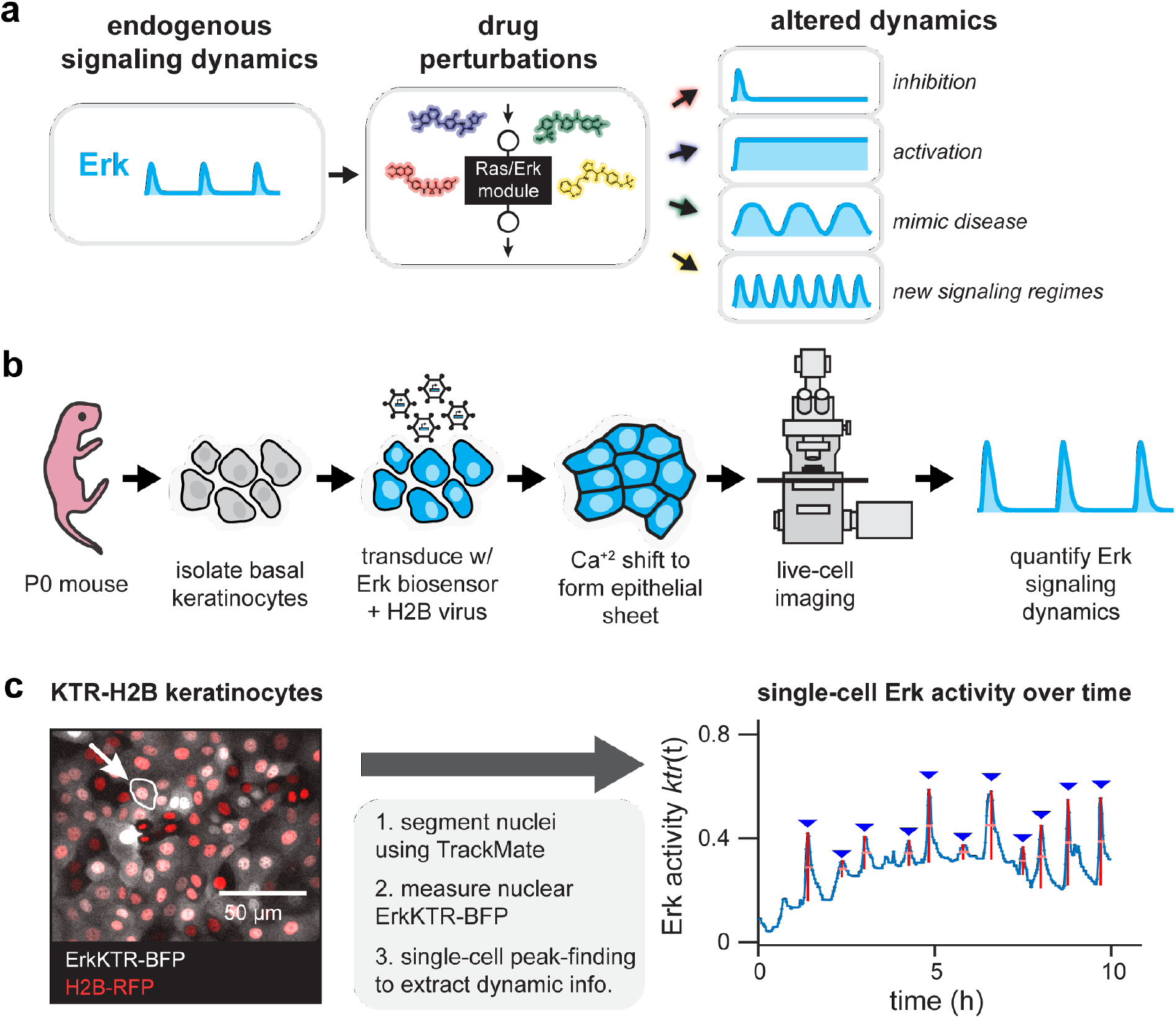
High-throughput screening for altered Erk dynamics. (A) A small-molecule screen for altered signaling dynamics. Such a screen can identify altered signaling behaviors including total pathway inhibition, constitutive activation, dynamics that resemble disease states, or previously undescribed dynamic regimes. (B) Primary keratinocytes as a model system for quantitative studies of Erk signaling dynamics. Basal epidermal keratinocytes are isolated from a P0 mouse pup, transduced with viral vectors encoding live-cell biosensors, induced to form a uniform epithelial sheet, and imaged via confocal microscopy. (C) An image processing pipeline for monitoring Erk dynamics. A fluorescent nuclear marker enables automated cell segmentation, tracking, and quantification of nuclear Erk kinase translocation reporter (KTR) activity. A representative image of KTR-BFP- and H2B-RFP-expressing primary mouse keratinocytes is shown, as well as quantification of Erk activity from the indicated cell.

Motivated by these possibilities, we set out to develop a screen for altered Erk signaling dynamics. Erk signaling is an ideal model system because of its stimulus-dependent dynamics (Marshall, 1995) and the excellent tools available for monitoring and controlling pathway dynamics in individual cells (Regot et al., 2014; Toettcher et al., 2013). We also report an excellent experimental model system for screening Erk dynamics: primary mouse keratinocytes, which we found to exhibit rapid, robust, and reproducible Erk dynamics even in the absence of externally-supplied growth factors. Screening a panel of 429 kinase inhibitors in more than 85,000 individual keratinocytes revealed multiple classes of altered Erk dynamics. The screen also uncovered previously-unknown regulatory links: inhibiting the non-EGFR receptor tyrosine kinases Met and VEGFR unexpectedly increased Erk’s steady-state activity and pulse frequency. Using optogenetic stimuli to directly control Ras/Erk dynamics confirmed that these drug-induced changes to dynamics are sufficient to alter cell proliferation. Our work thus begins to define the molecular processes that govern and regulate dynamic Erk activity, provides a compendium of drug-induced changes to signaling dynamics, and establishes an extensible experimental and computational platform for future live-cell screens.

## Results

### Primary keratinocytes are a robust model system for pulsatile Erk dynamics

We first needed to identify a cellular model of Erk dynamics that would be suitable for live-cell screening. Such a system must be highly dynamic, with frequent Erk pulses observed in a majority of cells. Dynamics should also be reproducible from day to day and robust to variations in conditions (e.g., media formulations; time post-plating). It was recently reported that Erk is highly dynamic in the basal epithelium of live adult mice (Hiratsuka et al., 2015), and epidermal keratinocytes can be cultured ex vivo without transformation or immortalization and are readily accessible to viral gene transduction and imaging (Nowak and Fuchs, 2009). We thus hypothesized that keratinocytes might serve as an ideal cell type to use as the basis for further dissection of Erk signaling dynamics.

We harvested basal keratinocytes from newborn CD1 mouse pups using established protocols (see **Methods**) (Nowak and Fuchs, 2009). After 10 passages, we transduced primary keratinocytes with viral vectors encoding a fluorescent biosensor of Erk kinase activity (KTR-BFP) and histone H2B (H2B-RFP) (**Figure 1B**) (Regot et al., 2014). We sorted these cells to obtain a homogenous population of dual-expressing, low-passage-number keratinocytes (termed KTR-H2B keratinocytes) that were used for all subsequent experiments. For analysis, we segmented and tracked H2B-RFP-positive nuclei, obtained nuclear KTR-BFP intensity for each cell, and quantified features of pulsatile Erk activity using the ImageJ plugin TrackMate and custom MATLAB code (**Figure 1C**; see **Methods** and **Supplementary Code, Figure S1**) (Abràmoff et al., 2004; Jaqaman et al., 2008; Tinevez et al., 2017).

We first set out to characterize baseline Erk activity dynamics in primary mouse keratinocytes. KTR-H2B keratinocytes were plated on fibronectin-coated glass, switched to high-calcium conditions to form an epithelial monolayer, and starved in growth factor-free (GF-free) media for 8 h prior to imaging. We observed frequent pulses of Erk activity in most cells, even though these conditions lacked any externally-supplied growth factors, serum, or other additives (**Figure 2A**, top; **Movie S1**). Quantifying Erk dynamics revealed that they were relatively stable over 24 h of imaging, with cells pulsing approximately once per hour on average (**Figure 2A**, bottom). Analysis of the timing between successive pulses revealed a broad distribution of waiting times with a long tail, suggesting that keratinocytes may exhibit stochastic, excitable Erk activity pulses rather than regular, periodic oscillations (**Figure S2A-B**). Similar Erk activity pulses were observed across a range of culture conditions, including skin-like organotypic cultures at an air-liquid interface (**Figure S2C-E**) (Aw et al., 2016). Immunostaining in embryonic mouse epidermis also revealed a salt-and-pepper pattern of Erk phosphorylation (**Figure S2F**), consistent with similar sporadic bouts of Erk activity in epidermal cells *in vivo* (Hiratsuka et al., 2015).

**Fig. 2.**
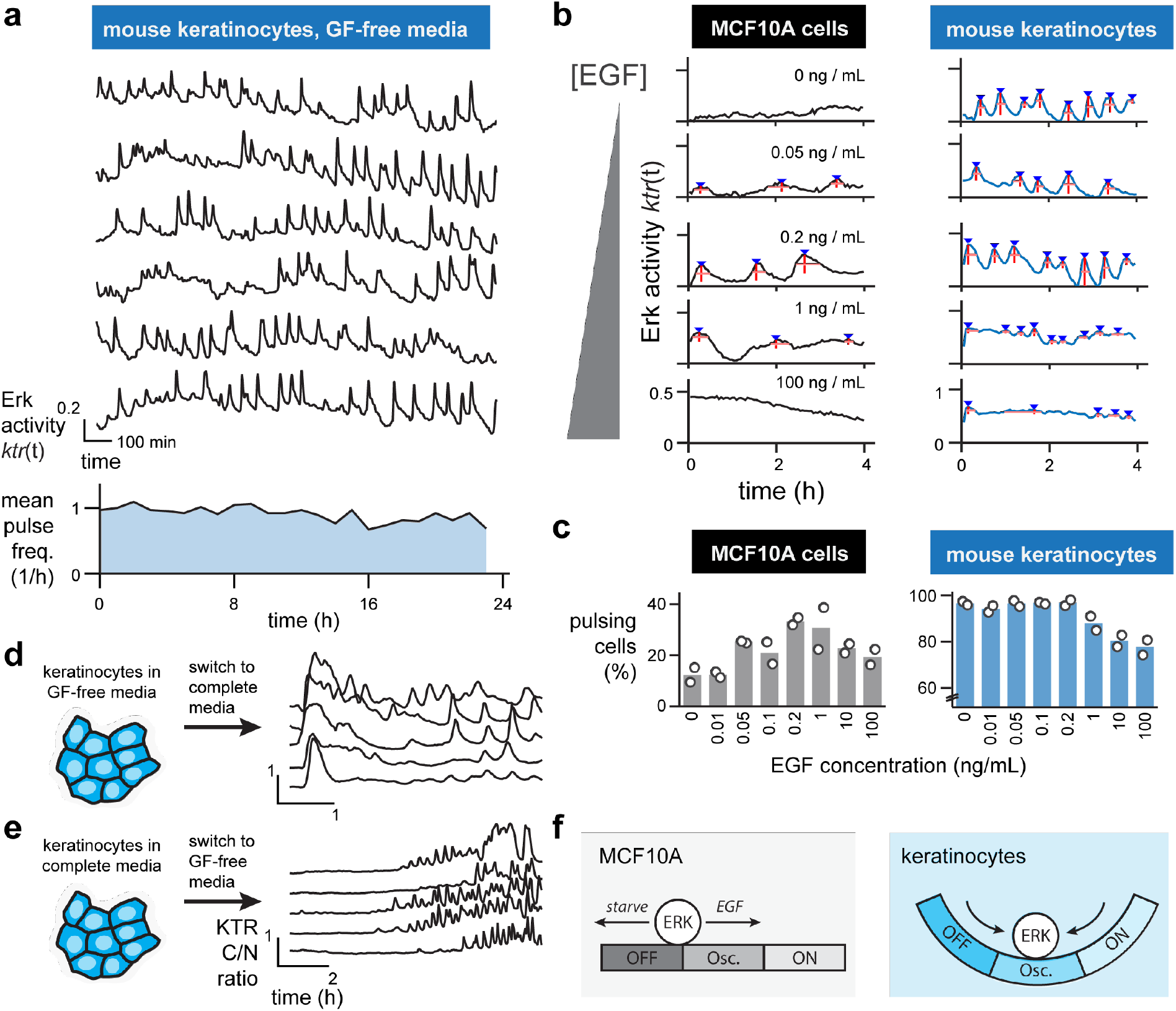
Primary keratinocytes exhibit an ‘idling motor’ ground state of pulsatile Erk dynamics. (**A**) Upper: representative single-cell traces of Erk activity in keratinocytes cultured in growth factor free (GF-free) media over 24 h. Lower: the mean number of pulses observed per cell for each hour over the 24 h timecourse. (**B**) Representative single-cell traces for MCF10A breast epithelial cells (left) and mouse keratinocytes (right) treated with increasing doses of epidermal growth factor (EGF). (**C**) The percentage of MCF10As (left) or mouse keratinocytes (right) that exhibited ≥ 2 Erk pulses over a 5 h timecourse after treatment with increasing doses of EGF. N=2 biological replicates (white circles); each replicate includes data from > 100 single cells. (**D-E**) Representative single-cell traces of Erk activity in keratinocytes shifted from GF-free to complete media (in **D**) and from complete to GF-free media (in **E**) showing the reappearance of pulsatile Erk dynamics over time. (**F**) Schematic comparing the stimulus-dependence of pulsatile Erk dynamics in classical cell lines such as MCF10As (left) and primary keratinocytes (right). Instead of a flat landscape where dynamics are permanently altered by GF stimulation, keratinocytes adapt and return to a ‘ground state’ of pulsatile Erk dynamics.

Unlike our observations in keratinocytes, prior studies in various epithelial cell lines found Erk dynamics to be strongly dependent on growth factors and culture conditions (Albeck et al., 2013; Aoki et al., 2013; Shankaran et al., 2009). We thus sought to directly compare keratinocytes to MCF10A epithelial cells, a classic model cell line for studying Erk dynamics (Albeck et al., 2013). We treated KTR-H2B-expressing MCF10As or keratinocytes with varying doses of epidermal growth factor (EGF) and monitored Erk activity for 4 h after stimulation (**Figure 2B; Figure S2G; Movie S2**). In MCF10As, Erk dynamics varied as a function of EGF concentration, from low, constant Erk activity in the absence of EGF to occasional pulses at intermediate EGF doses (50-200 pg/mL) and constant, high Erk activity in the presence of saturating EGF. Only 30% of MCF10A cells pulsed at least twice over 4 h of imaging, even at the most permissive EGF concentrations (**Figure 2C**, left). In contrast, keratinocytes exhibited robust, frequent Erk pulses across a broad range of EGF concentrations (from GF-free to 1 ng/mL EGF), with 80-100% of cells pulsing in all conditions tested (**Figure 2C**, right).

Our data led us to suspect that pulsatile Erk activity represents a baseline signaling state to which keratinocyte cells return, even after strong external perturbations. To test this model, we monitored Erk activity dynamics after acutely switching keratinocytes from GF-free to complete media or from complete to GF-free media (**Figure 2D-E**). Cells exhibited a high-amplitude peak of Erk activity in response to serum stimulation, followed by adaptation and a return to spontaneous pulsing over 2-3 h (**Figure 2D**). Conversely, switching to GF-free media drove Erk to a constant, low-activity state for 6 h, after which cells spontaneously resumed pulsing (**Figure 2E**). Primary mouse keratinocytes thus appear to harbor an ‘idling motor’ of pulsatile Erk signaling to which they return even after strong signaling perturbations (**Figure 2F**).

What might the function of these idling-motor Erk activity pulses be? One clue may lie in the ability of pulsatile, excitable systems to generate synchronous, high-amplitude responses to weak external stimuli, as has been suggested for cortical actin oscillations in *Dictyostelium* cells (Hoeller et al., 2016). Supporting this model, we observed that KTR-H2B keratinocytes exhibited a synchronous pulse of Erk activity when stimulated with 50 pg/mL EGF (**Figure S2H**), a dose 20 times lower than what is required to elevate cells’ overall Erk activity (1 ng/mL; **Figure S2I**). It is possible that this robust dynamic state may be a unique property of epidermal stem cells, perhaps related to their continuous lifelong proliferation. Alternatively, a robust Erk pulsatile state may be a universal property of mammalian epithelia *in vivo*, but has been lost over time in tissue-culture cell lines. A better understanding of *in vivo* Erk signaling dynamics will be crucial for discriminating these possibilities (Hiratsuka et al., 2015). Regardless, we concluded that keratinocytes’ ubiquitous and rapid Erk activity pulses make them an ideal model system to screen for compounds and pathways that regulate signaling dynamics.

### An imaging-based small molecule screen for altered Erk dynamics

We next set out to perform an imaging-based screen for compounds that alter Erk signaling dynamics. We selected a small molecule library consisting of 429 kinase inhibitors (Selleck Chemicals). Such a library is likely to be enriched for modulators of Erk dynamics, as kinase activity is a primary currency of MAPK signaling and crosstalk (Mendoza et al., 2011). Moreover, many kinase inhibitors have been developed to directly target EGFR/MAPK signaling and can be assessed as positive controls. Finally, increasingly comprehensive data on kinase inhibitor specificity is becoming available (Anastassiadis et al., 2011; Karaman et al., 2008; Moret et al., 2019) making it possible to move quickly from an initial hit to a candidate kinase target and potential molecular mechanism.

We developed an experimental approach to plate, starve, and stimulate KTR-H2B keratinocytes in 50 wells of a 384-well plate per imaging session (**Figure 3A**; see **Methods**). Precise volumes of each drug were delivered to a clean plate using an acoustic liquid handler, and drugs were then diluted to working concentrations in GF-free media and transferred to plates of keratinocytes to a 2.5 μM final concentration. Each batch of 50 wells included multiple vehicle-only (DMSO) controls to monitor batch-to-batch variability in Erk dynamics. We then performed two-color confocal imaging on 200 cells per drug every 3 min over 5 h, resulting in a total of 80,000 cells tracked and analyzed across all conditions.

**Fig. 3.**
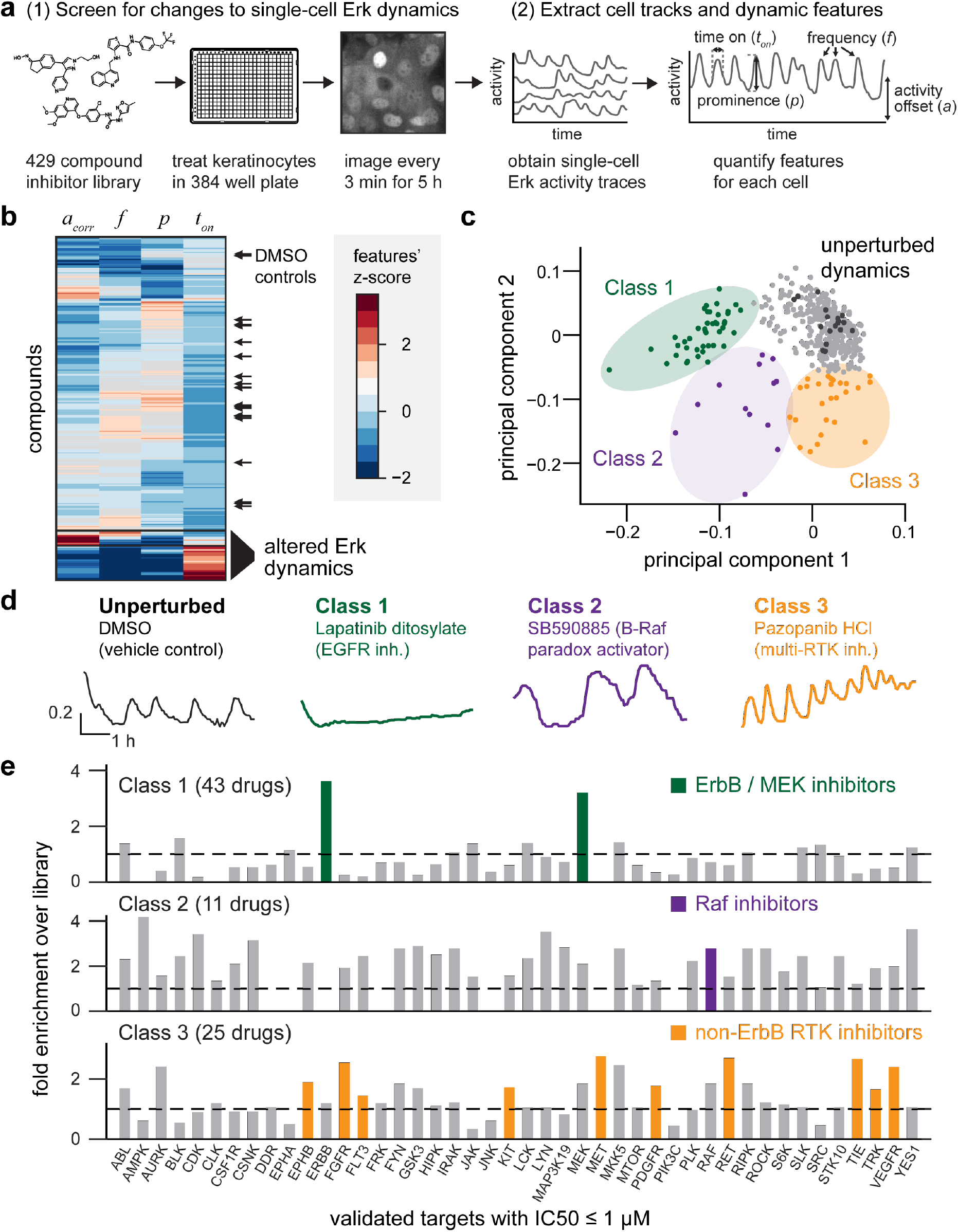
A kinase inhibitor screen for changes in Erk dynamics. (**A**) Overview of experimental and computational steps in the small molecule screen. KTR-H2B keratinocytes were plated in 384-well plates, treated with kinase inhibitors, and monitored every 3 min for 5 h. H2B and KTR fluorescence images were used to track cells, assess Erk activity, and extract dynamic features for approximately 200 single cells per well. (**B**) Hierarchical clustering of all compounds by four dynamic features. Black arrows: DMSO controls from all 9 plates. (**C**) Principal component analysis showing the contribution of each compound along the first two principal components. Black dots indicate DMSO controls, and three classes of drug-induced perturbations are indicated. (**D**) Representative single-cell traces of Erk dynamics for one drug from each class. Drug name and the expected molecular target are indicated. (**E**) Enrichment analysis for drug targets in each class. Enrichment scores represent how many compounds hit a target within a class as compared to the overall compound library. Highlighted bars indicate enriched targets representing specific biological hypotheses for each class.

We extracted three features of nuclear Erk dynamics for each cell – the mean pulse frequency *f*; the mean pulse prominence *p*; and the mean pulse width *t_on_* (**Figure S1**). We also measured the mean KTR cytosol-to-nuclear ratio at each timepoint to obtain the overall Erk activity offset, *a*, for each drug. Finally, we quantified the density of cells and their total displacement over the 5 h timecourse to test for indirect relationships between Erk dynamics and cell-physiological parameters. Importantly, a pairwise correlation analyses revealed that across all drugs, cell density was positively correlated with the Erk activity offset *a* (**Figure S3A-B**). Because plating density is an operator-controlled variable that is unrelated to which drug was added, we used a density-corrected Erk activity offset *a_corr_* for all analyses of the screen (**Figure S3C**).

Clustering by dynamic features revealed that most compounds did not alter Erk activity compared to DMSO controls, but that some drugs elicited strong changes in Erk dynamics (**Figure 3B**). To distinguish between different types of drug-induced dynamics, we performed principal component analysis on the matrix of compounds and dynamic measurements. This analysis revealed that the first two principal components (PCs) explained 88% of the dataset’s overall variance, with PC1 dominated by temporal features of the Erk pulses (pulse frequency *f* and pulse width *t_on_*), and PC2 was dominated by the amplitude features (activity offset *a* and pulse prominence *p*) (**Figure S3D**). Mapping each drug onto these principal components revealed three classes of drugs that were well-separated from the majority of compounds and DMSO controls (**Figure 3C; Table S1**). Supporting the association of PC1 and PC2 with Erk levels and timing, we found that plotting each compound’s activity offset and pulse frequency was also able to cleanly separate all three drug classes (**Figure S3E**).

The first class of 43 Erk-altering inhibitors (referred to here as ‘Class 1’) consisted of drugs that reduced Erk activity to a low, constant level, suggesting that this class may represent the canonical inhibitors of EGFR/MAPK signaling activity (**Figure 3D; Movie S3**). A second class of 11 drugs (referred to here as ‘Class 2’) elevated Erk activity and decreased pulse frequency (**Figure 3D; Movie S3**). Finally, a third class of 25 compounds (termed ‘Class 3’) amplified Erk activity by increasing the frequency and/or set point of Erk pulses (**Figure 3D; Movie S3**).

To gain insight into the kinases targeted by each drug class, we compiled available information about each drug’s molecular targets from Selleck Chemicals and the HMS LINCS project, a recently-developed database of experimentally-validated target information (Keenan et al., 2018; Moret et al., 2019). We then measured the enrichment of drug targets within each class over the entire library (**Figure 3E; Supplementary Methods**). This analysis was immediately useful in the case of Class 1 inhibitors, which was highly enriched for drugs with activity against EGFR-family (ErbB) receptors and the Erk-activating kinase MEK (**Figure 3E; Table S1**). Moreover, even Class 1 drugs that were not annotated as EGFR/MEK inhibitors (e.g., saracatinib, TWS119, ZM306416) have been implicated as direct inhibitors of these nodes in prior studies (Anastassiadis et al., 2011; Antczak et al., 2012; Formisano et al., 2015). Our drug screen thus correctly identified EGFR/MAPK pathway inhibitors as suppressors of dynamic Erk activity in keratinocytes. The observation that no other targets were enriched in this class also suggests that keratinocytes’ pulsatile Erk dynamics do not require permissive activity from other canonical kinase signaling pathways.

The limited number of Class 2 compounds precluded similarly informative enrichment analysis, as many kinases were only hit zero or one time (**Figure 3E**). However, examining individual drug responses revealed that two compounds in particular (SB590885 and GDC0879) elicited pronounced long, low-frequency Erk pulses (**Figure 3D**). Both compounds share a common classification as ‘paradox-activating’ B-Raf inhibitors that increase Raf activity by stabilizing Raf-Raf dimers (Hall-Jackson et al., 1999; Hatzivassiliou et al., 2010; Heidorn et al., 2010; Poulikakos et al., 2010). These results reinforce studies observing slow Erk dynamics in response to B-Raf mutation or paradox activation in other cellular contexts (Aoki et al., 2013; Bugaj et al., 2018). It is also notable that other paradox-activating B-Raf inhibitors in the compound library (e.g., vemurafenib) were not Class 2 hits. This may be explained by the observation that SB590885 is an especially potent inducer of Raf dimerization at micromolar concentrations, unlike other paradox-activating inhibitors including vemurafenib (Miyamoto and Sawa, 2019). There may thus be a spectrum of paradoxical Raf activation, with changes in Erk dynamics reflecting the degree to which Raf dimers are stabilized.

We finally turned our attention to Class 3 drugs, which elicited high-frequency Erk pulses on an elevated baseline (**Figure 3D**). Enrichment analysis revealed that no single kinase target was highly enriched; instead, we observed a broad enrichment for compounds that target many receptor tyrosine kinases *other* than EGFR, including VEGFR, Met, Kit, and FGFR (**Figure 3E**). Conversely, Class 1 was depleted for inhibitors targeting these same non-EGFR RTKs (**Figure 3E**). The observation that EGFR inhibitors and non-EGFR RTK inhibitors have opposing effects on Erk activity in keratinocytes led us to hypothesize that various receptor tyrosine kinases may exert antagonistic, inhibitory crosstalk on EGFR-driven Erk pulses. Such RTK-level crosstalk may provide a previously unappreciated layer of regulation over Erk dynamics, and we sought to better delineate its role in the following sections.

### Inhibiting non-EGFR receptor tyrosine kinases tunes EGFR-triggered Erk dynamics

Our drug screen suggested that inhibitors of various receptor tyrosine kinases might unexpectedly activate Erk, and we next sought to validate this class of targets and confirm their dynamics-altering effects. We focused on Met and VEGFR, receptor tyrosine kinases whose inhibitors were especially potent for increasing Erk pulse frequency, and which have each been shown to be expressed in keratinocytes and to play functional roles in epidermal tissue (Chmielowiec et al., 2007; Wilgus et al., 2005). We repeated drug stimulus experiments with fresh aliquots of tivozanib (targeting VEGFR), pazopanib (VEGFR), cabozantinib (Met + VEGFR), and the Class 2 compound GDC0879 (B-Raf), confirming that each drug was sufficient to alter Erk dynamics (**Figure 4A; Movie S4**).

**Fig. 4.**
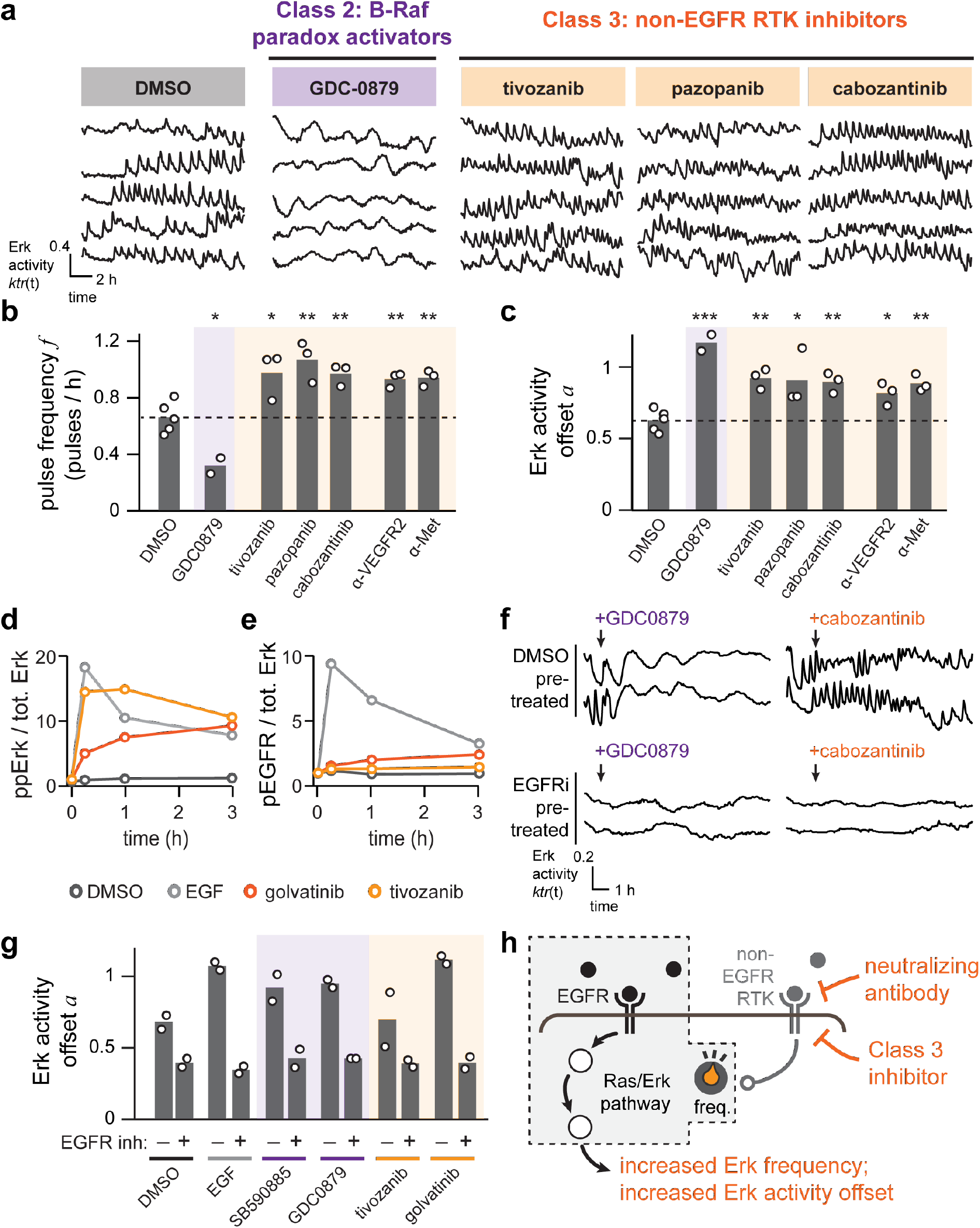
Inhibiting non-EGFR receptor tyrosine kinases increases dynamic Erk activity in keratinocytes. (**A**) Representative traces from keratinocytes treated with either a Class 2 B-Raf inhibitor, a Class 3 RTK inhibitor, or DMSO as a control. Cells were imaged every 3 min over 12 h. (**B-C**) Average Erk pulse frequencies (in **B**) and overall Erk activity offset (in **C**) for keratinocytes treated as in **A** or with a neutralizing antibody targeting Met or VEGFR2. Each biological replicate (white circle) represents dynamics assessed from at least 100 single cells. Statistics are derived using a two-sided t-test (*, p<0.05; **, p < 0.01; ***, p < 0.001). (**D-E**) Western blot quantification for phospho-Erk (ppErk; in **D**) and phospho-EGFR (pEGFR; in **E**) in keratinocytes after the addition of either EGF or a non-EGFR RTK inhibitor (golvatinib or tivozanib). pEGFR and ppErk are normalized to total Erk protein and each value is represented as fold-induction relative to the untreated initial timepoint. (**F**) Representative single-cell traces showing Erk activity in keratinocytes that were pre-treated for 1 h with the EGFR inhibitor lapatinib or DMSO control, and then stimulated with EGF, Class 2 or 3 inhibitors, or additional DMSO. (**G**) The overall Erk activity offset a for cells treated as in **F**. Each biological replicate (white circle) represents the mean of at least 100 cells imaged for 8 h. (**H**) A conceptual model summarizing the connection between RTK inhibition and increasing Erk activation. Inhibition of non-EGFR receptor tyrosine kinases by small molecules or neutralizing antibodies acts downstream of EGFR to enhance EGFR-driven pulsatile Erk signaling.

We next sought to confirm that Met and VEGFR are indeed the relevant molecular targets responsible for the change in Erk dynamics. We thus tested whether a separate class of inhibitors, neutralizing antibodies against RTKs’ ligand-binding domains, could elicit similar responses. We treated KTR-H2B keratinocytes with either Met or VEGFR neutralizing antibodies, kinase inhibitors, or vehicle controls, and monitored Erk dynamics over time. Class 3 kinase inhibitors and VEGFR/Met neutralizing antibodies each elicited significant changes in Erk frequency and activity offset (**Figure 4B-C**). Together, these results demonstrate that inhibiting non-EGFR RTK activity indeed modulates Erk dynamics.

How might inhibition of non-EGFR RTKs alter activity within the EGFR/Ras/Erk network? One possible nexus of crosstalk is the EGF receptor itself, as prior studies of cancer therapeutics indicate that inhibiting one family of RTKs can induce the activation of parallel ligand-receptor pairs (Brand et al., 2014; Engelman et al., 2007). To test whether Met and VEGFR inhibition directly modulates EGFR activity, we used immunoblotting to measure EGFR and Erk phosphorylation at various timepoints after addition of the VEGFR inhibitor tivozanib or the Met/VEGFR inhibitor golvatinib (**Figure 4D-E; Figure S4**). We observed high levels of Erk phosphorylation in response to both drugs, matching the increase in baseline Erk activity observed for Class 3 compounds in the drug screen (**Figure 4D**). In contrast, EGFR phosphorylation was unaffected by these compounds (**Figure 4E**), indicating that non-EGFR RTK inhibitors enhance Erk activity without altering the activity state of EGFR.

Our observations so far are consistent with two models. First, a compound might act to increase MAPK activity directly, functioning independently of EGFR-mediated activation. Alternatively, a compound may modulate or amplify signaling generated by the intact EGFR/MAPK network. The latter model would be expected in the case of Class 2 drugs, as it is known that paradox-activating Raf inhibitors depend on upstream Ras activity for enhanced signaling (Heidorn et al., 2010). To distinguish between these possibilities, we monitored Erk dynamics after addition of both an EGFR inhibitor and an Erk-modulating compound (**Figure 4F-G**). We found that in all cases, EGFR inhibitor pre-treatment blocked the Erk-stimulatory effect of Class 2 and 3 drugs. These data support a model where inhibition of non-EGFR receptor tyrosine kinases alters transmission through the intact EGFR/MAPK signaling pathway, tuning Erk’s pulse frequency and overall activity level (**Figure 4H**).

### Drug-induced changes to Erk signaling dynamics have distinct effects on cell proliferation

A growing body of literature suggests that signaling dynamics can be interpreted into specific cell fate outcomes (Johnson and Toettcher, 2019; Marshall, 1995; Purvis and Lahav, 2013). Yet in most cases, which dynamic features are sensed by cells to trigger a response is still unknown. We thus wondered whether the Erk dynamics observed in our pharmacological screen might also produce distinct cellular responses. In addition to revealing principles of cell fate control, such differences may be relevant to clinical responses after RTK inhibitor treatment. We focused on cell proliferation as a model Erk-driven cellular response (**Figure 5A**), as prior work has demonstrated that Erk activity is sufficient for proliferation under a range of conditions (Albeck et al., 2013; Aoki et al., 2013; Bugaj et al., 2018; Toettcher et al., 2013).

**Fig. 5.**
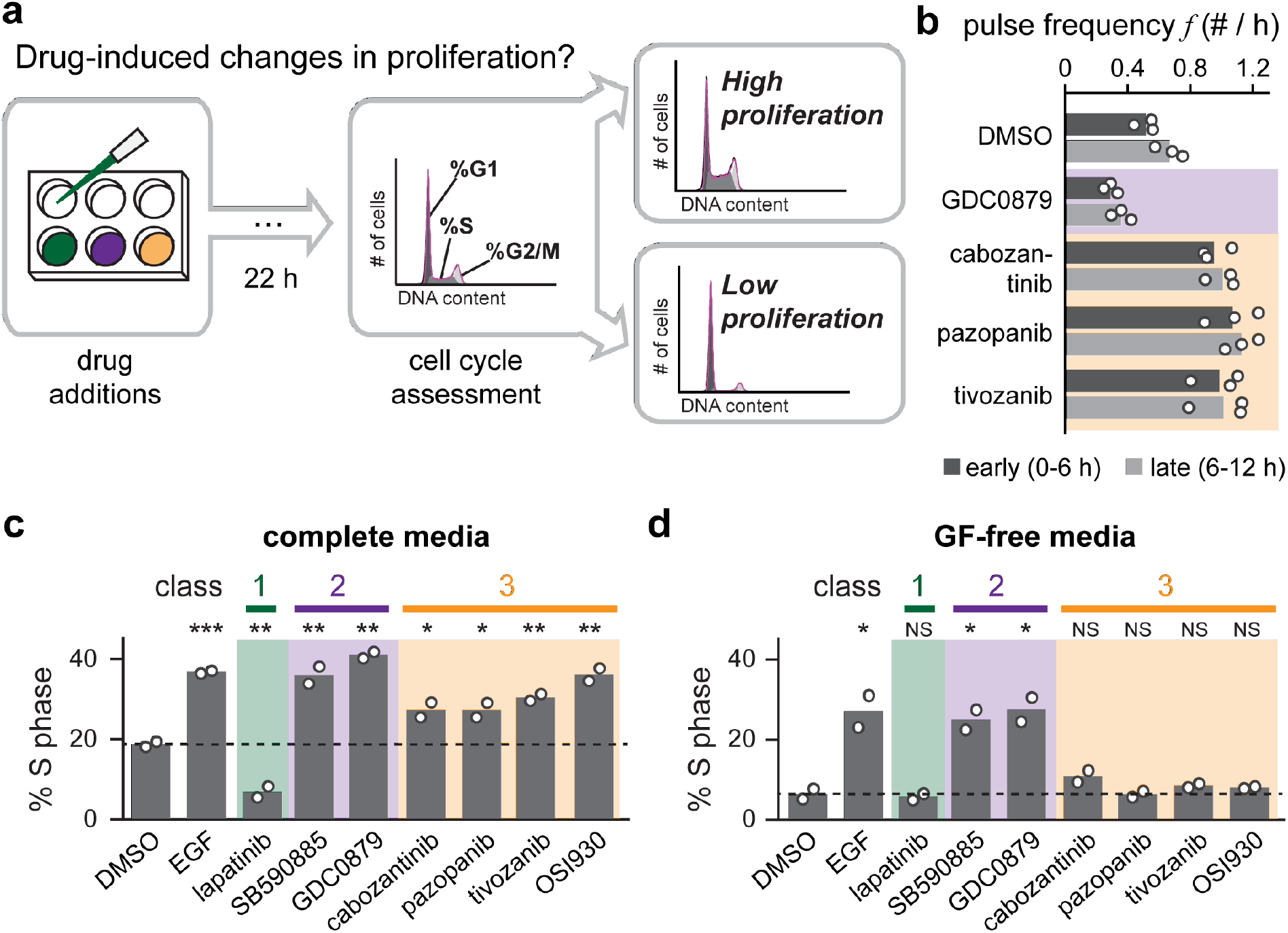
Drugs that alter Erk dynamics have distinct effects on cell-proliferative responses. (**A**) Schematic describing assay for cell proliferation in response to drug treatment. Keratinocytes are incubated for 22 h in the presence of a candidate compound and the resulting cell cycle distribution is assessed by flow cytometry. The fraction of S phase cells is taken as a proxy for proliferation. (**B**) Quantification of Erk activity pulse frequency during the first and second halves of 14 h timecourses after stimulation with Class 2 and 3 compounds. Each biological replicate (white circle) represents the mean of least 100 single-cell trajectories. (**C-D**) The three classes of dynamics-altering drugs produce distinct proliferative responses. The percentage of cells in S phase was measured for keratinocytes treated with compounds from Class 1 (green), Class (purple), and Class 3 (orange), EGF, or DMSO control in complete media (in **C**) or GF-free media (in **D**). Biological replicates (white circles) were assessed from flow cytometry DNA content distributions of at least 20,000 cells each. Statistics are derived using a one-sided t-test (*, p < 0.05; **, p < 0.01; ***, p < 0.001).

Mammalian cells’ commitment to proliferation is thought to occur over an 8-12 h window, culminating in the onset of S-phase (Zetterberg and Larsson, 1985; Zwang et al., 2011). We thus first set out to validate that drug-induced changes to Erk dynamics were persistent over at least the 8-12 h required for cell cycle entry. Using previously-collected 14 h imaging data of Erk dynamics in response to compounds from each class (**Figure 4A**), we quantified dynamics separately in the first and second halves of the dataset. We observed similar drug-induced shifts to pulse frequency in both the first and second 7-hour period (**Figure 5B**), indicating that drug-induced changes to dynamics are indeed sufficiently persistent to impact proliferation.

We next set out to assess drug-induced changes in cell proliferation. We first cultured keratinocytes in complete media, a condition that is permissive for proliferation but where drug-induced changes in Erk dynamics can still be observed (**Figure S5**). We then applied compounds from each of the three drug classes and measured the percentage of S-phase cells after 22 h (**Figure 5A**). We found that the Class 1 inhibitor lapatinib strongly reduced proliferation, consistent with a requirement for Erk activity to drive cell proliferation (**Figure 5C; Figure S6A**). In contrast, Class 2 and 3 compounds that increased overall Erk activity also enhanced proliferation, increasing the proportion of S-phase cells comparable to a saturating dose of EGF.

Repeating the same experiment in GF-free media revealed similar responses in two cases: Class 1 compounds suppressed proliferation, whereas Class 2 compounds increased it. However, although Class 3 compounds elicited high levels of proliferation in complete media, they were unable to drive cell proliferation in GF-free media (**Figure 5D; Figure S6B**), even though they produce fast, frequent Erk pulses in both cases. These data raised the possibility that the not all Erk-activating stimuli are interpreted equally, and that the fast dynamics elicited by Class 3 drugs may trigger proliferation under a more restrictive set of conditions than the slow dynamics elicited by Class 2 drugs.

### Optogenetic stimuli causally link Erk dynamics to cell proliferation

So far, we have seen that certain classes of kinase inhibitors alter both Erk dynamics and cell proliferation, but this correlative relationship must be interpreted with caution. Each drug may alter multiple features of Erk dynamics (e.g., pulse frequency; overall Erk activity) and likely acts on multiple kinase targets in the cell (Giuliano et al., 2018), making it impossible to unambiguously assign a dynamic response to a specific cellular outcome. We thus set out to directly control Erk dynamics and assess their effects on cell proliferation using an optogenetic strategy (Bugaj et al., 2018). We transduced keratinocytes with a blue-light-sensitive variant of the OptoSOS optogenetic system (Toettcher et al., 2013) that we have found to function robustly in contexts ranging from mammalian cells to *Drosophila* embryos (Guntas et al., 2015; Johnson et al., 2017).

Our goal is to completely control Erk dynamics with light, yet keratinocytes exhibit endogenous Erk pulses that could play a confounding role. We reasoned that pre-treatment with the Class 1 drug lapatinib could be used to block endogenous EGFR-induced pulses, while still enabling us to deliver light inputs to Ras and activate Erk signaling on demand (**Figure 6A**). If successful, such a system would enable us to vary the frequency of light pulses to mimic the shifts in Erk dynamics that are elicited by Class 2 and 3 drugs (**Figure 6B**). To validate this strategy, we monitored Erk dynamics in KTR-H2B-OptoSOS keratinocytes that were treated with lapatinib and toggled between darkness and blue light every 15 min. We observed consistent, controllable Erk dynamics for more than 8 h (**Figure 6C; Movie S5**). Light stimuli can thus be used to precisely control Erk signaling even in cells that normally harbor complex, autonomous EGFR-driven signaling dynamics.

**Fig. 6.**
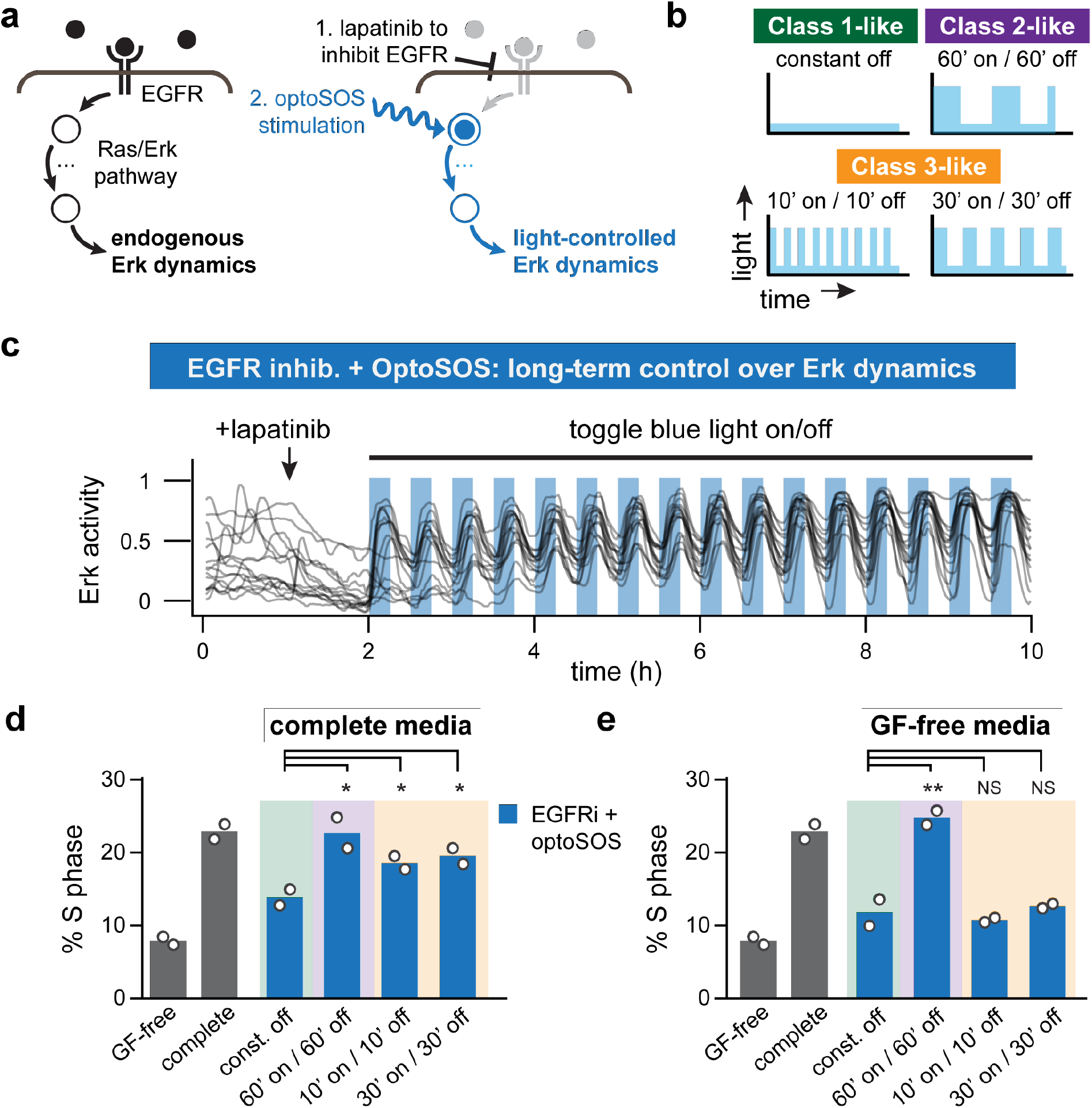
Direct optogenetic control over Erk dynamics reveals drug-like proliferative responses. (**A**) Schematic of our strategy to implement direct optogenetic control in keratinocytes, which requires overriding their endogenous dynamics and establishing light-controlled Erk activity pulses. Endogenous Erk dynamics are first eliminated by treating cells with an EGFR inhibitor, then the OptoSOS system is used to apply light-dependent optogenetic inputs directly to Ras. (**B**) Light pulses designed to mimic each class of drug-induced Erk dynamics: Class 1-like suppression of Erk activity, Class 2-like low-frequency pulses, or Class 3-like high-frequency pulses. (**C**) Representative single-cell Erk activity traces from keratinocytes stimulated as in A. Cells’ endogenous dynamics were imaged for 1 h, then the EGFR inhibitor lapatinib was added to extinguish endogenous Erk signaling, after which blue light was toggled on and off every 15 min. (**D-E**) Proliferation in response to light-driven Erk dynamics. Blue bars indicate OptoSOS keratinocytes grown in complete media (in **D**) or GF-free media (in **E**), treated with the EGFR inhibitor lapatinib (blue bars), and then subjected to repeated blue light pulses at the schedules indicated. Gray bars indicate control conditions: continuous GF-free media or complete media, both without lapatinib or light stimulation. For all conditions, S phase entry was assessed after 22 h. Biological replicates (white circles) were assessed from flow cytometry DNA content distributions of at least 20,000 cells each. Statistics are derived using a one-sided t-test (*, p < 0.05; **, p < 0.01; ***, p < 0.001).

We next set out to measure proliferation in response to Erk activity dynamics that mimic the effects of Class 2 and Class 3 drugs. A primary difference between these two drug classes is their shift in the frequency of Erk pulses. To isolate the role of this frequency shift, we designed light inputs that maintained the same overall dose of blue light but were pulsed at different frequencies (10’on/10’off, 30’on/30’off, or 60’on/60’off). Pulses were delivered using an Arduino microcontroller driving LED light panels in a tissue-culture incubator, and DNA content was assessed after 22 h (see **Methods**). We found that the proliferative responses triggered by optogenetic Ras/Erk pulses broadly matched those obtained previously from Class 1, 2, and 3 compounds: both high-frequency and low-frequency light pulses partially restored proliferation in lapatinib-treated cells cultured in complete media (**Figure 6D; Figure S7A**), just as had been observed for Class 2 and Class 3 inhibitors (**Figure 5C**). Optogenetic stimulation was also able to replicate the context-dependence of high-frequency Erk pulses, which failed to induce proliferation in GF-free media (**Figure 6E; Figure S7B**), just as had been observed for Class 3 drugs (**Figure 5D**). Finally, 60’on/60’off pulses, reminiscent of the dynamics induced by paradox-activating Raf compounds, increased the proliferation of lapatinib-treated cells in both complete and GF-free media (**Figure 6E**). These results confirm that Erk dynamics are sufficient to explain each of the effects on cell proliferation that we observed in our pharmacological classes, including the inability of high-frequency Erk pulses to drive proliferation in GF-free media.

## Discussion

### Primary keratinocytes as a platform for studying Erk signaling dynamics

Many signaling pathways exhibit complex, time-varying patterns of activity (Purvis and Lahav, 2013), yet in most cases it is unknown how these dynamics are generated, regulated, and interpreted. Small-molecule and genetic screens could be powerful tools to better understand dynamics (Behar et al., 2013), as they have provided fundamental insights into the origins of complex phenotypes for over a century (Morgan, 1910; Nusslein-Volhard and Wieschaus, 1980). Yet the field is only beginning to define how signaling dynamics screens should be designed and interpreted. One challenge is that dynamics can span many timescales, with pulses of activity that may rise or fall within minutes but occur only a few times per day (Albeck et al., 2013; Lahav et al., 2004; Nandagopal et al., 2018), necessitating minutes-timescale imaging over multi-day time periods. Dynamics can also be highly variable between cells and experimental conditions (Albeck et al., 2013; Lahav et al., 2004), requiring automated tracking and analysis of large numbers of cells to accurately estimate dynamic features.

The current study provides one path towards a successful signaling dynamics screen. A crucial factor was the discovery that primary keratinocytes harbor rapid and robust Erk dynamics, enabling us to observe multiple pulses from nearly all cells within 5 h (**Figure 2**) in a manner that was reproducible from day to day (**Figure 3**). A second facilitating factor involved combining recent advances in automated nuclear segmentation and tracking (Jaqaman et al., 2008; Tinevez et al., 2017) and an Erk biosensor whose activity can be quantified simply from measurements of nuclear fluorescence (Regot et al., 2014). We argue that the combination of excellent biosensors, image processing tools, and an appropriate model system makes Erk dynamics an ideal test bed for future high-throughput studies of signaling dynamics.

The dynamics we observe in keratinocytes begin to constrain potential models of how Erk pulses are generated. We observed high-amplitude Erk pulses with a broad range of time intervals between successive pulses in unstimulated keratinocytes (**Figure S2A-B**), as well as Erk pulses that can be synchronized by low-dose EGF stimulation (**Figure S2I**). These data suggest that keratinocytes harbor an excitable pulse generator, analogous to the sporadic spikes of membrane potential during neuron firing (Hodgkin and Huxley, 1952). Moreover, some Class 3 drugs further increased the frequency of Erk pulses up to one every 20 min (Figure 4A), a time-scale that is likely to be too fast to be driven by cycles of protein expression/degradation or ligand secretion/removal. Rather, our data suggest pulsatile Erk activity that is driven by a network of protein-protein interactions and post-translational modifications (Rust et al., 2007).

The current study is only a first step toward comprehensive high-throughput analysis of intracellular signaling dynamics. Our experimental design (450 experimental conditions analyzed over nine 5-hour imaging sessions) is far from an upper limit and could be increased by using a higher well density and a large field-of-view camera to capture multiple wells in a single image, reducing imaging time per condition, and increasing the total number of imaging sessions. We anticipate that these improvements could increase throughput by 1-2 orders of magnitude, sufficient to screen 10,000 conditions and opening the door to genome-wide RNAi or CRISPR approaches (Wang et al., 2014). Merging live-cell imaging with genome-wide perturbation screening could profoundly transform our understanding of the molecular mechanisms driving intracellular signaling dynamics.

### Inhibiting RTKs can paradoxically increase Erk activity and pulse frequency

Our small-molecule screen revealed distinct ways in which kinase inhibitors can alter Erk pathway activity (**Figure 7A**). One immediate, intuitive result was that Class 1 inhibitors confirmed a classic model of MAPK signaling: inhibiting EGFR, MEK, or Erk is sufficient to block pulsatile Erk signaling for hours, and no other families of kinase inhibitors exhibit a similarly suppressive effect. The screen also offered some surprises beyond this classic view. For instance, few kinases are known to naturally suppress MAPK signaling such that their inhibition would result in increased Erk activity. Yet our screen identified almost as many inhibitors that increased Erk activity as those that suppressed it (36 vs. 43, respectively). A closer look at these Erk-activating hits could thus uncover new regulatory inputs into a canonical cell signaling pathway.

**Fig. 7.**
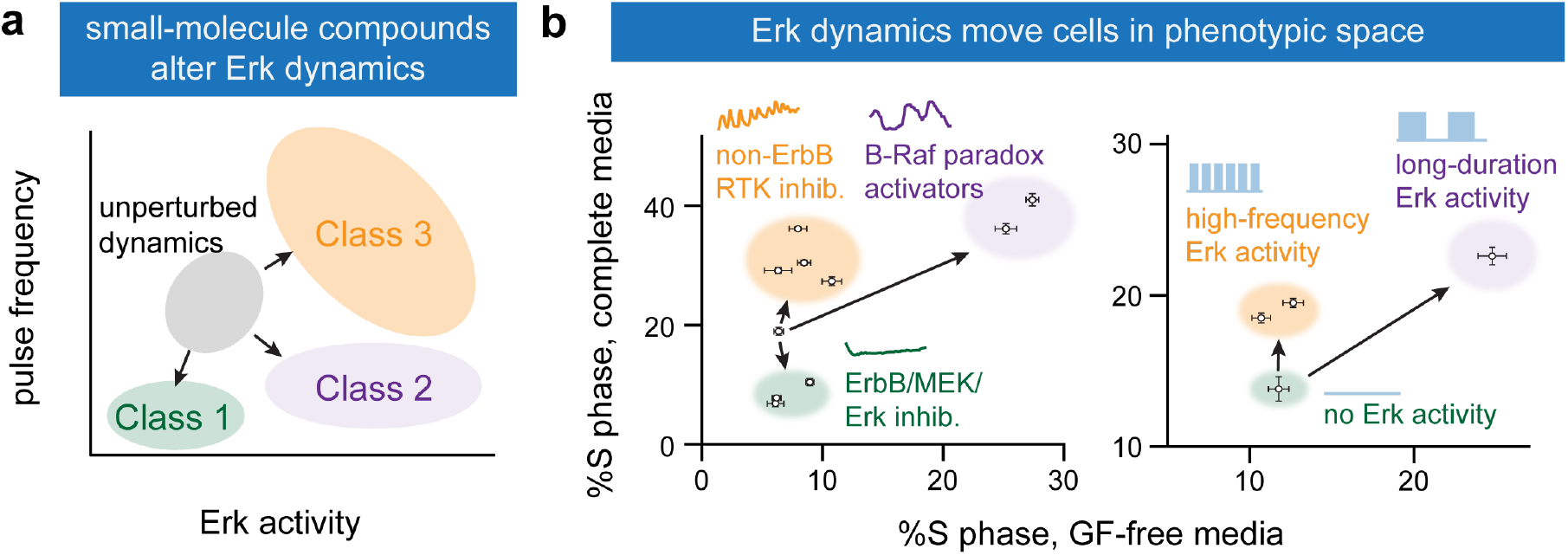
Insights into signaling dynamics and cell fate control revealed by a small molecule screen and optogenetics. (**A**) Compounds from a small-molecule screen alter Erk signaling dynamics along two dimensions. Class 1 compounds block Erk activity, Class 2 inhibitors drive long, slow pulses, and Class 3 inhibitors increase Erk pulse frequency on an elevated background. (**B**) Changes in Erk dynamics have functional consequences for the cell. Cell proliferation in different environments represent a single point in ‘phenotypic space’. Two perturbations that alter Erk dynamics, small molecule treatment (left) and optogenetic control (right), shift populations of cells to different positions in phenotypic space. Blocking Erk activity inhibits proliferation in all conditions (green); increasing Erk pulse frequency enhances proliferation in complete media but not GF-free media (yellow); long, slow Erk pulses enhance proliferation in both complete and GF-free media. Points (mean ± SEM) were obtained from the experimental data presented in **Figure 5C-D** and **Figure 6D-E**.

One class of Erk-activating inhibitors has been the subject of considerable recent interest: paradox-activating B-Raf inhibitors (Heidorn et al., 2010; Poulikakos et al., 2010). In our screen, two Class 2 drugs of this type dramatically increased overall Erk activity and slowed its dynamics, extending the duration of pulses from 10 min to over 1 h. The slow dynamics that we observe in primary epidermal keratinocytes also match recent reports in human lung epithelial cell lines (Bugaj et al., 2018) and the MDCK canine kidney cell line (Aoki et al., 2013), suggesting that modes of perturbation to Erk dynamics may apply broadly across many cellular contexts.

Our study also defined an additional, unreported class of Erk-activating compounds targeting non-EGFR receptor tyrosine kinases, which elicited fast, high-frequency Erk pulses on an elevated baseline. As in the case of B-Raf, this observation runs counter to expectations because RTKs are canonically viewed as activators of Erk. It should be noted that reactivation of Erk signaling through parallel receptor/ligand combinations is an important mechanism of cancer cells’ resistance to RTK inhibition (Bean et al., 2007; Dong et al., 2011; Engelman et al., 2007; Zhang et al., 2012). Yet this model of drug-induced resistance also fails to explain our data; we observe a shift in Erk dynamics immediately after drug addition (**Figure 4F**), suggesting that it is not a product of network rewiring caused by long-term drug exposure.

How might non-EGFR RTK inhibition increase Erk activity? We find that inhibitor-triggered Erk activation requires continued EGFR signaling, but that EGFR phosphorylation is unaffected (**Figure 4D-G**). These data suggest that inhibiting non-EGFR RTKs somehow potentiates signaling through an intracellular node between EGFR and Erk. Although a mechanistic picture is still unclear, it is tempting to speculate that RTKs might compete for access to a limiting pool of intracellular scaffold proteins and effectors (e.g., Grb2, Shc, SOS). In this model, inhibiting one receptor tyrosine kinase would increase the effector pool available to transduce another receptor’s signaling output. Pulsatile signaling may thus enable two receptors to alternate access to a limiting pool of scaffold proteins, allowing multiple RTKs to ‘share’ intracellular signaling channels like the Ras/Erk pathway.

We identified Met and VEGFR as two receptors whose inhibition increases Erk activity (**Figure 4**). One might logically question whether receptors that bind to ligands named ‘hepatocyte growth factor’ (HGF, the ligand for Met) and ‘vascular endothelial growth factor’ (for VEGFR) play functional roles in the basal epidermis. Not only do epidermal keratinocytes express Met (Sennett et al., 2015), but HGF is known to promote their migration, proliferation, and survival (Busch et al., 2008; Locatelli and Lange, 2011) and Met signaling in keratinocytes is critical for wound healing (Chmielowiec et al., 2007). Similarly, all VEGFR isoforms are expressed by epidermal keratinocytes (Man et al., 2006) and have also been shown to drive keratinocyte proliferation, migration, and wound healing (Man et al., 2008; Wilgus et al., 2005; Wu et al., 2014). These broadly overlapping biological roles and similar effects on Erk dynamics (Figure 4A) suggest that one or more regulatory connections are shared by both receptors to tune EGFR/Erk activity in epidermal cells. Importantly, our data also raise the possibility that a broad range of clinical RTK inhibitors may unexpectedly act to increase EGFR/MAPK pathway activity in the skin.

### Differences in Erk dynamics can drive proliferative decision-making

The small-molecule screen revealed how endogenous Erk signaling dynamics could be pharmacologically altered, but what can these perturbations teach us about how dynamics are functionally interpreted? To address this question, we examined a classic Erk-driven response – cell proliferation – in complete and GF-free media. Cells’ behavior in these two environments can be thought of as points in a two-dimensional ‘phenotype space’. Unperturbed keratinocytes occupy one point in this space, as they proliferate in complete but not in GF-free media (**Figure 7B**). Using both optogenetic and pharmacological perturbations, we found that Erk dynamics moved cells to three distinct positions in this phenotype space. Blocking Erk activity reduced proliferation (**Figure 7B**; green region), whereas long, slow Erk pulses increased proliferation in all cases (**Figure 7B**; purple region). In contrast, short Erk pulses, even delivered at a high frequency, were only able to increase proliferation in complete media and elicited no response in GF-free conditions (**Figure 7B**; orange region).

Our data reveal that the mapping between Erk activity and cell proliferation is more complex than is typically thought. The total dose of Erk activity is not simply interpreted into a defined proliferative outcome (Johnson and Toettcher, 2019) nor does proliferation require continuous, persistent Erk activity in keratinocytes (Zetterberg and Larsson, 1985). Instead it appears that the duration of pulses may be directly sensed and, in the case of high-frequency stimuli, interpreted with additional serum-induced cues to determine the cell’s response (Zwang et al., 2011). This complex combinatorial logic was unexpected at the outset of our study, yet it is likely to be central to two seemingly-contradictory observations: that keratinocytes maintain robust Erk pulses in GF-free media and yet are unable to proliferate in these conditions. Our data resolves this contradiction: we find that frequent, brief pulses of Erk activity are insufficient for proliferation on their own, and additional GF-dependent signals or longer-duration pulses are necessary to mount a proliferative response. We eagerly await future studies to reveal the molecular ‘decoding circuits’ that enable dynamic and combinatorial intracellular signals to be translated into a cellular response such as S-phase entry (Jeknic et al., 2019).

## ACKNOWLEDGEMENTS

We thank all members of the Toettcher and Devenport labs for their insights and comments. This work was supported by NIH Ruth L. Kirschstein NRSA fellowship F30CA206408 (to A.G.G.) and by NIH grant DP2EB024247 and NSF CAREER Award 1750663 (to J.E.T.). We also thank Dr. Christina DeCoste and the Princeton Flow Cytometry Resource Center for flow cytometry support; Dr. Gary Laevsky and the Princeton Molecular Biology Microscopy Core, which is a Nikon Center of Excellence, for microscopy support; and Hahn Kim and the Princeton Small Molecule Screening Facility for support during the compound screen.

## Methods

### Cell culture

Dorsal epidermal keratinocytes derived from CD1 mice and stably expressing a retrovirally-delivered histone H2B-RFP (selected for expression with hygromycin) were obtained from the Devenport lab (B. Heck) and were cultured as described previously (Nowak and Fuchs, 2009). Briefly, keratinocytes were given low calcium (50 μM) growth media (E media supplemented with 15% serum and 0.05 mM Ca2+) in Nunc Cell Culture Treated Flasks with filter caps (Thermo) and were maintained in a humidified incubator at 37° C with 5% CO2. Cell passage number was kept below 30.

### Plasmids

The pHR backbone (Addgene 79121) was used for all lentiviral expression constructs (Zufferey et al., 1998). All lentiviral constructs were cloned using PCR amplification of complementary fragments and In-Fusion HD-based assembly (Takara). ErkKTR-BFP was cloned by PCR amplification of TagBFP (Addgene 102350) and insertion into the pHR ErkKTR-iRFP construct used previously (Addgene 111510). BFP-SSPB-SOScat-2A-PuroR-2A-iLID-CAAX was cloned from the pTiger-OptoSOS construct used previously (Addgene 86439), followed by PCR amplification and insertion of 2A-PuroR (a gift from Brett Stringer; Addgene 98290) and TagBFP. ErkKTR-iRFP-2A-H2B-tRFP was cloned by PCR amplification of the 2A cleavage site from pTiger-OptoSOS and amplification of the H2B-tagRFP coding sequence (a gift from Sergi Regot; Addgene 99271), followed by insertion into pHR ErkKTR-iRFP. H2B-RFP retrovirus was previously constructed in the Devenport lab (B. Heck) using the pSINRevPGK-hyg backbone (S. Williams, Elaine Fuchs lab).

### Lentivirus production and stable transduction

Lentivirus was generated as described previously (Goglia et al., 2017). Briefly, Lenti-X HEK293T cells were seeded at roughly 50% confluency in 6-well dishes. Cells were allowed to settle for 12 h, after which FuGENE HD transfection reagent was used to co-transfect cells with the desired pHR expression vector and the two necessary lentivirus helper plasmids (pCMV-dR8.91 and pMD2.G – gifts from the Trono lab). Viral supernatants were collected 48-52 h after transfection, cell debris was removed using a 0.45 μm filter, and viruses were either used immediately or stored at −80° C.

For viral transduction, keratinocytes were plated at <50% confluency in 6-well dishes, allowed to adhere overnight, and then treated with 100 μL of the desired lentivirus. Viral supernatants were supplemented with 5 μg/mL Polybrene and 50 μM HEPES buffer. Virus-containing media was removed after 24 h and cells were transferred into new tissue culture flasks. Fluorescence-activated cell sorting (FACS) was used to isolate keratinocytes that stably expressed both high levels of the H2B-RFP nuclear marker and low levels of the ErkKTR-BFP reporter. Sorting was performed using a FACSAriaIII Fusion system with 355, 405, 488, 561, and 637 nm laser excitation sources (BD Biosciences). For optogenetic experiments, a separate line of OptoSOS keratinocytes was generated using the same lentiviral transduction procedures outlined above and then, 3d after transduction, selecting for expression by exposing cells to puromycin (2 μg/mL) for > 3d. Sorted/selected cells were then expanded into multiple large culture flasks and 20 vials of each sorted line were frozen down and stored in liquid nitrogen for use during the study.

### Preparing cells for microscopy

Imaging experiments were performed in either 384- or 96-well black-walled, 0.17mm high performance glass-bottom plates (Cellvis). Before plating, the bottom of each well was pre-treated with a solution of 10 μg/mL bovine plasma fibronectin (Thermo Fisher) in phosphate buffered saline (PBS) to support cell adherence. Two days before imaging, keratinocytes were seeded at 16,000 cells/well in 50 μL of low-calcium E media (in a 384-well plate), plates were briefly centrifuged at 100 × g to ensure even plating distribution, and cells were allowed to adhere overnight. 24 h before imaging, wells were washed 2-3X with PBS to remove non-adherent cells and were shifted to high-calcium (1.5 mM CaCl2) complete E media to promote epithelial monolayer formation. For experiments in GF-free media, cells were washed once with PBS and shifted to high-calcium P media (i.e., DMEM/F12 containing only pH buffer, penicillin/streptomycin, and 1.5 mM CaCl2) eight hours before imaging. To prevent evaporation during time-lapse imaging, a 50 μL layer of mineral oil was added to the top of each well immediately before imaging.

The kinase inhibitor library was screened in rounds of 48 drugs plus two DMSO controls per plate for a total of 50 wells, as this was the maximum number of conditions that could be imaged every 3 min in two fluorescent channels. Thus, nine initial rounds of imaging were required to screen all 429 drugs in the library. Cell plating was staggered such that a fresh plate of 50 wells in GF-free media was ready to be imaged every 8 h. An additional two rounds were added to re-screen drugs from wells with insufficient cell density.

### Microscopy

Imaging was performed on a Nikon Eclipse Ti confocal microscope, with a Yokogawa CSU-X1 spinning disk, a Prior Proscan III motorized stage, an Agilent MLC 400B laser launch containing 405, 488, 561, and 650 nm lasers, a cooled iXon DU897 EMCCD camera, and fitted with an environmental chamber to ensure cells were kept at a 37° C and 5% CO2 during imaging. All images were captured with either a 10X or 20X air objective and were collected at intervals of 2-3 min.

### Image processing and quantitative analysis of Erk dynamics

#### Processing raw TIFF files into tracked/segmented Erk activity dynamics

Our analysis pipeline takes as input a two-color, multi-timepoint TIFF stack that includes two fluorescent channels – KTR-BFP and H2B-RFP – for monitoring Erk activity and tracking/segmenting nuclei. These TIFF images contained metadata including the spatial scale of the data, and since our analysis pipeline incorporates true spatial units (μm rather than pixels), it can be used for images collected with any objective magnification. We first used a FIJI Jython script `batch_trackmate.py` to automatically run the TrackMate segmentation/tracking plugin (Jaqaman et al., 2008; Tinevez et al., 2017) on a folder of TIFF stacks, returning the mean nuclear intensity for each cell in both the H2B and KTR channels. Each dataset thus consisted of two paired folders: one containing all the TIFFs (with names <image_name>.tif) and one of the corresponding TrackMate tracks (with names <image_name>.txt). In addition to containing the mean nuclear intensities in all fluorescent channels, the resulting TrackMate tracks also include the X-Y locations of each nucleus at all timepoints, enabling analysis of each cell’s overall displacement and velocity.

#### Analyzing Erk activity dynamics from tracked cells

All subsequent analyses were performed in MATLAB. We wrote a set of analysis tools to automatically load the TIFFs and TrackMate tracks from the paired folders and analyze them with a set of options specified in a comma-delimited parameters file that is generated for each dataset. We first exclude TrackMate-tracked nuclei on either of two conditions: (1) they do not express both H2B and KTR, or (2) if H2B intensity changes dramatically over the imaging timecourse, indicating a tracking error (**Figure S1A-B**). We then filter and process nuclear KTR intensities using the following steps: (1) data is background-subtracted; (2) data is h-minima transformed to ‘flatten’ noise between consecutive timepoints; (3) data is inverted and scaled assuming that the maximum nuclear KTR intensity observed for each cell corresponds to Erk activity of 0; and (4) data is fed into a peak-finding routine that splits the trajectory into windows containing individual peaks, which are then analyzed using MATLAB’s built-in `findpeaks.m` function (**Figure S1C-D**).

#### Estimating the overall Erk activity offset for the entire well

In addition to analysis of single-cell trajectories, we also perform one well-averaged measurement at each timepoint: the overall cytosol-to-nuclear ratio computed for all identified cells. To do so, we used image dilation and erosion operations to obtain an annulus of cytoplasm surrounding all segmented nuclei. We then computed the mean C/N ratio by measuring the mean intensity in all cytosolic pixels divided by the mean intensity of all nuclear pixels (**Figure S1E-F**). We found this ratio to be an excellent proxy for well-averaged activity; as an example, it is able to accurately detect keratinocytes’ graded Erk responses to increasing doses of EGF (**Figure S1G**). We opted for a well-averaged C/N ratio rather than a single-cell ratio primarily because correctly assigning cytosolic regions to the appropriate nucleus can be quite tricky, especially in an epithelial monolayer.

At the conclusion of our analysis routine, the user is provided with two data structures: well, which contains the raw and processed nuclear KTR and Erk activity trajectories; and PS, a structure that contains all analyses of Erk dynamics over time (e.g., frequency, pulse prominence, overall Erk activity offset, total distance moved per cell, etc.). These quantities were used for all subsequent analyses. Example runs are available on Github (github.com/toettchlab/Goglia2019).

### Optogenetics experiments

For microscopy experiments involving optogenetic stimuli, an X-Cite XLED1 light source coupled to a Polygon400 Mightex Systems digital micromirror device (DMD) was used to stimulate cells with spatially and temporally precise patterns of 450 nm blue light, applied for 500 ms every 1 min.

For proliferation experiments involving optogenetic stimuli, light was delivered using custom-printed circuit boards of blue 450 nm light-emitting diodes (LEDs). During light stimulation, cells were maintained in an incubator at 37° C in separate foil-wrapped boxes covered with separate blue LED boards delivering different patterns of light inputs. Each LED board was connected to a separate constant-current LED driver, all of which were controlled using an Arduino MEGA 2560 microcontroller board. The Arduino was programmed with open-source IDE software to deliver different dynamic light input regimes to each circuit board. To minimize phototoxicity, light inputs were delivered in cycles of 10 sec ON and 20 sec OFF – this allowed us to minimize light exposure while still delivering a constant stimulus to cells by taking advantage of the slow (0.5-1 min) dark decay rate of iLID activation (Guntas et al., 2015).

### Drug treatments

#### Drug screen

An Echo^®^ acoustic liquid handler (Labcyte) was used to precisely spot plastic 96-well plates with 75 nL of either DMSO vehicle control or of individual drugs from the kinase inhibitor library (Selleck Chem). 100 μL of P media was then added to each well that had been spotted by the Echo to create 3X stock solutions that could be rapidly added to plates of cells by multi-channel pipetting. Keratinocytes were plated on glass 384-well dishes in 50 μL of media and a final drug concentration of 2.5 μM was achieved by adding 25 μL of each 3X stock solution to individual wells of keratinocytes. Imaging was carried out 30 min after drug additions.

#### Subsequent testing of Class 1, 2, and 3 hits

After downstream analyses of the primary screen, new aliquots of all top drug hits were purchased from Selleck Chemicals and drug effects were confirmed via extended time-lapse imaging. For all subsequent experiments involving confirmed drug hits (e.g., individual drug follow-ups, proliferation assays), drug additions were performed by first creating a 10X working stock by diluting drug/growth factor/neutralizing antibody in GF-free media and then adding an appropriate volume of this stock to cultured cells.

#### Determining EGFR-dependence of Class 3 drug effects

KTR-H2B keratinocytes were prepared for imaging in multiwell plates as before. Prior to imaging, half the wells were pre-treated with the EGFR inhibitor lapatinib at 2.5 μM while the other half were given a DMSO vehicle control and Erk dynamics were monitored by time-lapse confocal microscopy. After 1 h of imaging, representative Class 2 and Class 3 compounds were added to either DMSO- or lapatinib-pre-treated cells and the resulting Erk dynamics were monitored over 8 h.

### Cell lysate collection and Western blotting

Two days before lysate collection, cells were seeded at 3 × 106 in 6-well tissue culture dishes and allowed to adhere overnight. 24 h before collection, cells were washed 3X with PBS and shifted to high-calcium media to promote epithelial monolayer formation. Drugs/growth factors were added to cells at the indicated times before lysate collection (all drugs added at 2.5 μM; EGF added at 10 ng/mL). At indicated time points, media was aspirated, cells were washed with PBS and were lysed with 100 μL of ice-cold RPPA buffer (1% Triton X-100, 50 mM HEPES buffer, 150 mM NaCl, 1.5 mM MgCl2, 1 mM EGTA, 100 mM NaF, 10 mM sodium pyrophosphate, 1 mM Na3VO4, 10% glycerol, freshly-prepared protease/phosphatase inhibitors). Cell scrapers were used to remove cells from the surface of each well, then each lysate was transferred to an ice-cold Eppendorf tube and centrifuged at 17,000 × g for 10 min at 4° C. Supernatants were transferred to new tubes, 25 μL of 4X NuPAGE LDS Sample Buffer (Thermo Fisher) was added to each, and samples were boiled at 98° C for 5 min. Samples were then stored at −80° C.

For Western blotting, lysates were run in 17-well 4-12% Bis-Tris gels (Thermo Fisher) at 120V for 1.5 h. A Trans-Blot SD semi-dry transfer cell (Bio-Rad) was used to transfer protein samples from gels to nitrocellulose membranes. Single-gel transfers were run at constant current of 70mA and max voltage of 25V for 1 h. Nitrocellulose membranes were then blocked for 1 h at room temperature in Odyssey blocking buffer (Li-Cor) and were incubated overnight at 4° C on a plate rocker in a 1:1000 dilution of primary antibody in blocking buffer. The following antibodies were used: anti-phospho-Erk1/2 rabbit monoclonal antibody (Cell Signaling 4370), anti-phospho-EGFR (pY1068) rabbit monoclonal antibody (Cell Signaling 3777), and anti-total-Erk1/2 mouse monoclonal antibody (Cell Signaling 4696). The next day, blots were washed 5 × 5 min with TBST and then incubated for 1 h at room temperature in a 1:10,000 dilution of IRDye 680RD goat anti-mouse and 800CW goat anti-rabbit fluorescent secondary antibodies (Li-Cor). Blots were then washed 5 × 5 min with TBST, imaged on a Li-Cor Odyssey CLx imaging system, and images were analyzed using Image Studio software (Li-Cor).

### Immunofluorescence staining

E14.5 embryos were dissected in PBS and fixed with 4% paraformaldehyde for 1 h. Dissected back skins were washed 3 × 5 min with PBS and then placed in permeabilization / blocking buffer (0.3% Triton X-100, 2% normal goat serum, 2% normal donkey serum, 2% bovine serum albumin, and 1% fish gelatin in PBS) for 2 h at room temperature. Samples were then incubated overnight on a plate rocker at 4° C in a 1:200 dilution of anti-phospho-Erk1/2 (Thr202/Tyr204) rabbit monoclonal antibody (Cell Signaling 4370) in blocking buffer. The next morning, samples were washed 3 × 30 min with blocking buffer and then incubated for 2 h at room temperature in a 1:750 dilution of Alexa Fluor 488-conjugated donkey anti-rabbit secondary antibody (Thermo Fisher A21206) and 1 μg/mL DAPI in blocking buffer. Tissues were again washed 3 × 30 min with blocking buffer, mounted on coverslips in Fluoro-Gel mounting medium (Electron Microscopy Sciences, Cat. 17985-30), and imaged by confocal microscopy.

### Organotypic keratinocyte culture in air-liquid interface

500,000 primary KTR-H2B keratinocytes were seeded on inverted 1.1 cm2 hanging cell culture inserts fit with 0.4 μm pore polyethylene terephthalate (PET) filters (Millipore PIHT15R48) in high calcium E media (1.5 mM). For the first 3 days, cells were grown with media on either side of the filter and were provided with fresh media daily. An air-liquid interface (ALI) was established on day three by removing all media, inverting the cell culture insert to hang in a 24-well dish, and then replacing media only inside the cell culture insert (i.e., removing media from above the cells) (**Figure S2F**). Cells were grown in ALI for an additional 3 days and were provided with fresh media daily. 6-day cultures were inverted, placed on cover slips and live-imaged with a spinning disk confocal microscope in a temperature- and CO2-controlled incubation chamber.

### Proliferation and cell cycle analysis

Cells were plated as described for western blotting above and were incubated with appropriate drug/growth factor/optogenetic stimuli for 22 h before analysis. Cells were then trypsinized, spun down, and resuspended in PBS containing 0.1% Triton X-100, 200 μg/mL propidium iodide (Sigma), and 200 μg/mL RNAse A (Sigma) to lightly permeabilize and stain for DNA content. Samples were incubated in the dark for 20-30 min and analyzed by flow cytometry (Darzynkiewicz and Juan, 2001). 20,000 single cells were analyzed for each experimental replicate. All flow cytometry was performed on a BD LRSII Multi-Laser Analyzer using a 561 nm laser. Cell cycle fractions were determined using FlowJo V.X software by Watson (Pragmatic) modeling.

### Statistical analyses

Statistical significance was determined using a standard t-test for p-values. For **Figure 4**, p-values were calculated using a two-sided t-test assuming equal variances because we sought to test for both positive and negative effects on signaling dynamics by small molecules. For **Figures 5–6**, p-values were calculated using a one-sided t-test assuming equal variances because we specifically tested the hypothesis that each treatment would alter cell proliferation in proportion to its effect on Erk dynamics (e.g., increased Erk -> increased proliferation).

### Data and software availability

All Jython and MATLAB code is available on Github (github.com/toettchlab/Goglia2019). All time-lapse microscopy data from the small-molecule screen will be available at the Image Data Resource (idr.openmicroscopy.org/; accession number forthcoming).

