## Supplementary Information for "A live-cell screen for altered Erk dynamics reveals principles of proliferative control"

**This PDF file includes:**

Supplementary Figures 1-7
Supplementary Tables 1 and 2
Supplementary Video 1-5 Legends

### Supplementary Figures and Legends

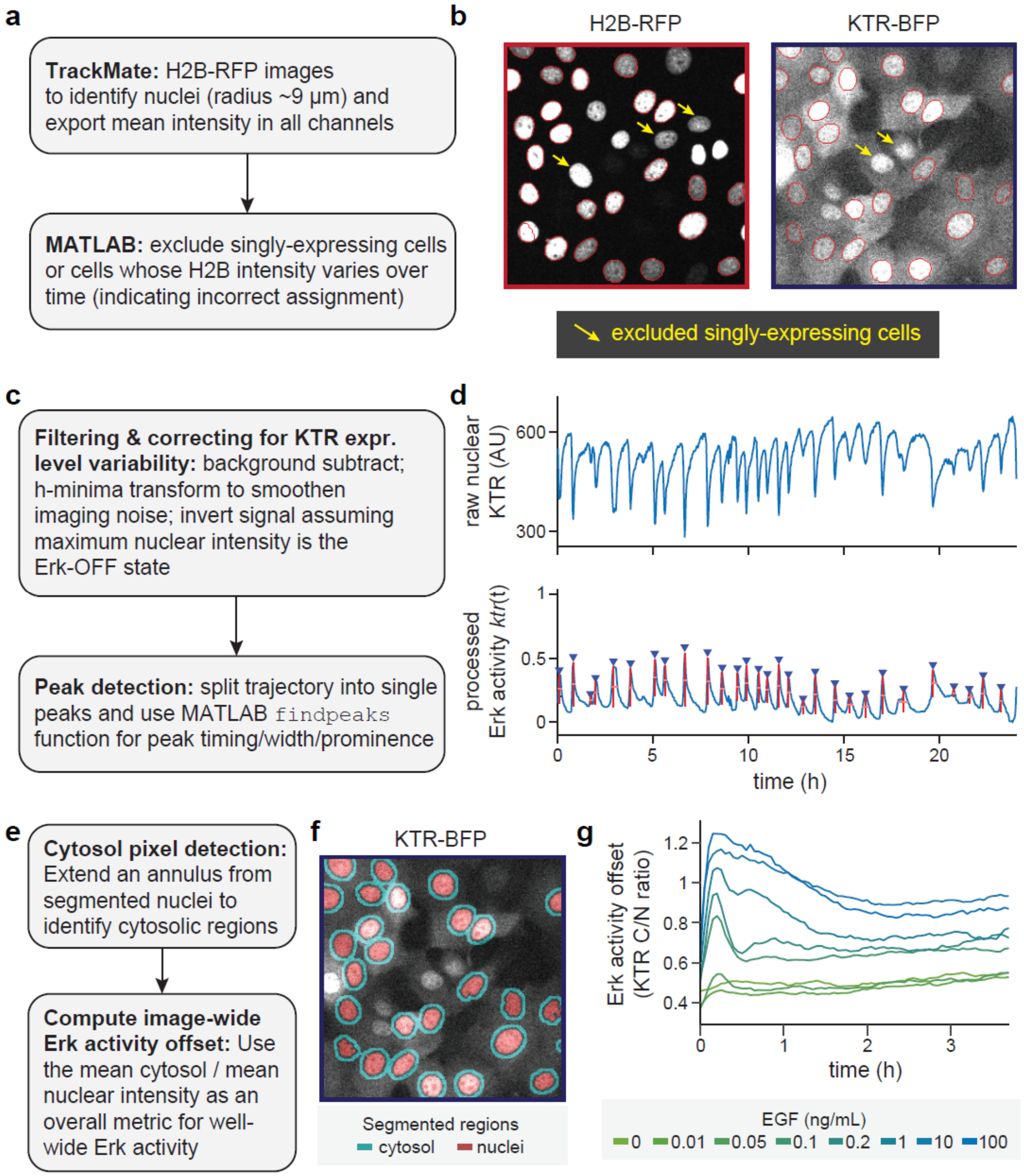

**Figure S1, related to Figure 1. Data processing pipeline for microscopy data of Erk activity.** (**A,C,E**) Schematics of each component of the data processing pipeline that takes in multi-channel TIFF image stacks of H2B-RFP and KTR-BFP as inputs, and returns quantitative data about single-cell Erk dynamics. (**B**) Images of H2B and KTR images where nuclei have been automatically identified (red circles). Nuclei are excluded that only express one of the two components (yellow arrows). (**D**) Filtering and processing of Erk activity trajectories. Representative trajectory of nuclear KTR fluorescence from a single keratinocyte over time (upper panel) is h-minima transformed, inverted, and subjected to peak-finding (lower panel). Erk activity pulse timing (blue triangles), pulse prominence (red vertical bars), and pulse width (pink horizontal bars) are obtained for each trajectory. (**F**) For quantifying well-averaged Erk activity, cytosolic regions are computed by a combination of dilation and erosion operations to define regions outside of nuclei. The ratio of the mean cytosolic and nuclear pixel intensities is used as a quantitative metric of the overall Erk activity offset at each timepoint. (**G**) Quantifying Erk activity offset for wells of keratinocytes stimulated with EGF demonstrates that a pulsatile Erk response can be sensitively detected for EGF concentrations as low as 50 pg/mL.

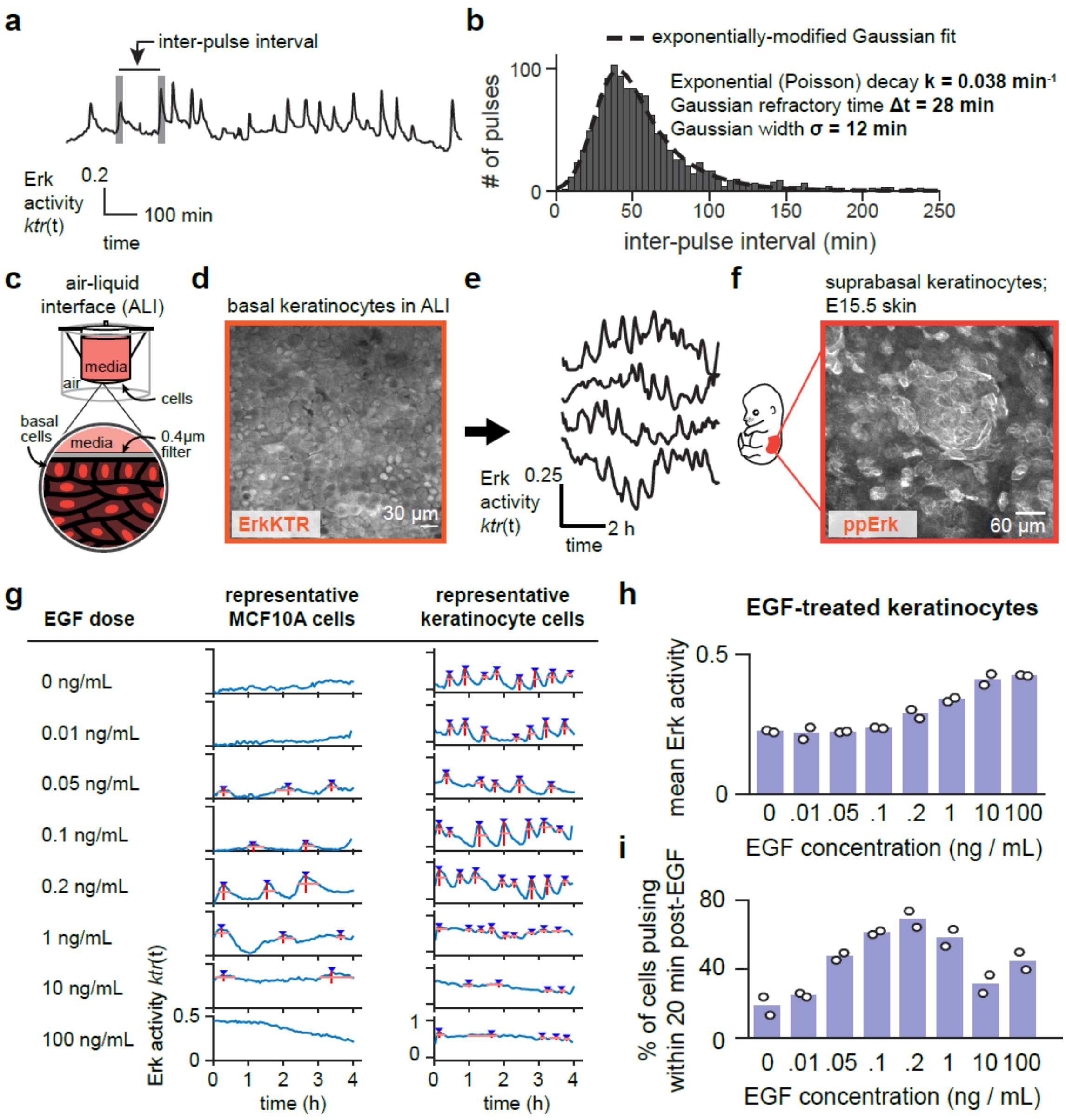
a
inter-pulse interval
Erk activity ktr(t)
0.2
100 min
time
b
exponentially-modified Gaussian fit
Exponential (Poisson) decay k = 0.038 min-1
Gaussian refractory time Δt = 28 min
Gaussian width σ = 12 min
### of pulses
inter-pulse interval (min)
c
air-liquid interface (ALI)
media
air
cells
basal cells
0.4μm filter
d
basal keratinocytes in ALI
ErkKTR
30 μm
e
Erk activity ktr(t)
0.25
2 h
time
f
suprabasal keratinocytes; E15.5 skin
ppErk
60 μm
g
EGF dose
representative MCF10A cells
representative keratinocyte cells
0 ng/mL
0.01 ng/mL
0.05 ng/mL
0.1 ng/mL
0.2 ng/mL
1 ng/mL
10 ng/mL
100 ng/mL
Erk activity ktr(t)
time (h)
h
EGF-treated keratinocytes
mean Erk activity
EGF concentration (ng / mL)
i
% of cells pulsing within 20 min post-EGF
EGF concentration (ng / mL)

**Figure S2, related to Figure 2. Comparing and quantifying Erk dynamics in keratinocytes and MCF10A cells**. (**A-B**) Quantifying the distribution of 'inter-pulse intervals' (depicted in **A**) can be used to characterize the extent of periodic vs stochastic pulses. A tight Gaussian distribution at a single value is indicative of a periodic system; an exponentially-decaying tail indicates stochastic pulses firing at a characteristic rate. An admixture of periodic and stochastic behavior would generate a Gaussian-exponential distribution (in **B**). The distribution of intervals between each Erk pulse over a 24 h timecourse is depicted here (gray bars), fit to an exponentially-modified Gaussian distribution (dotted black line). Parameters of the fit indicate the exponential decay time and Gaussian time between periodic pulses. (**C**) Schematic of organotypic air-liquid interface (ALI) keratinocyte culture (see **Methods**). Cells are cultured on a hanging insert exposed to media on their basal surface. (**D**) Confocal image of the basal layer of KTR-H2B keratinocytes grown as an ALI for 6 d. (**E**) Representative Erk activity trajectories for cells in **D** imaged over time. (**F**) Images of whole-mount epidermis taken from E15.5 mouse embryos and immunostained for phospho-Erk1/2 (See **Methods**). (**G**) Representative Erk activity traces and identified pulses from single MCF10A (left) or primary keratinocyte (right) cells treated with increasing doses of EGF. (**H-I**) Average Erk activity offset *a* (in **H**) and the fraction of cells pulsing within 20 min of EGF treatment (in **I**) for keratinocytes treated with the indicated dose of EGF. A maximal pulsatile response is observed at 200 pg/mL of EGF, whereas maximum overall Erk activity is observed at 10 ng/mL. Each biological replicate (white circle) represents dynamics assessed from at least 100 single cells.

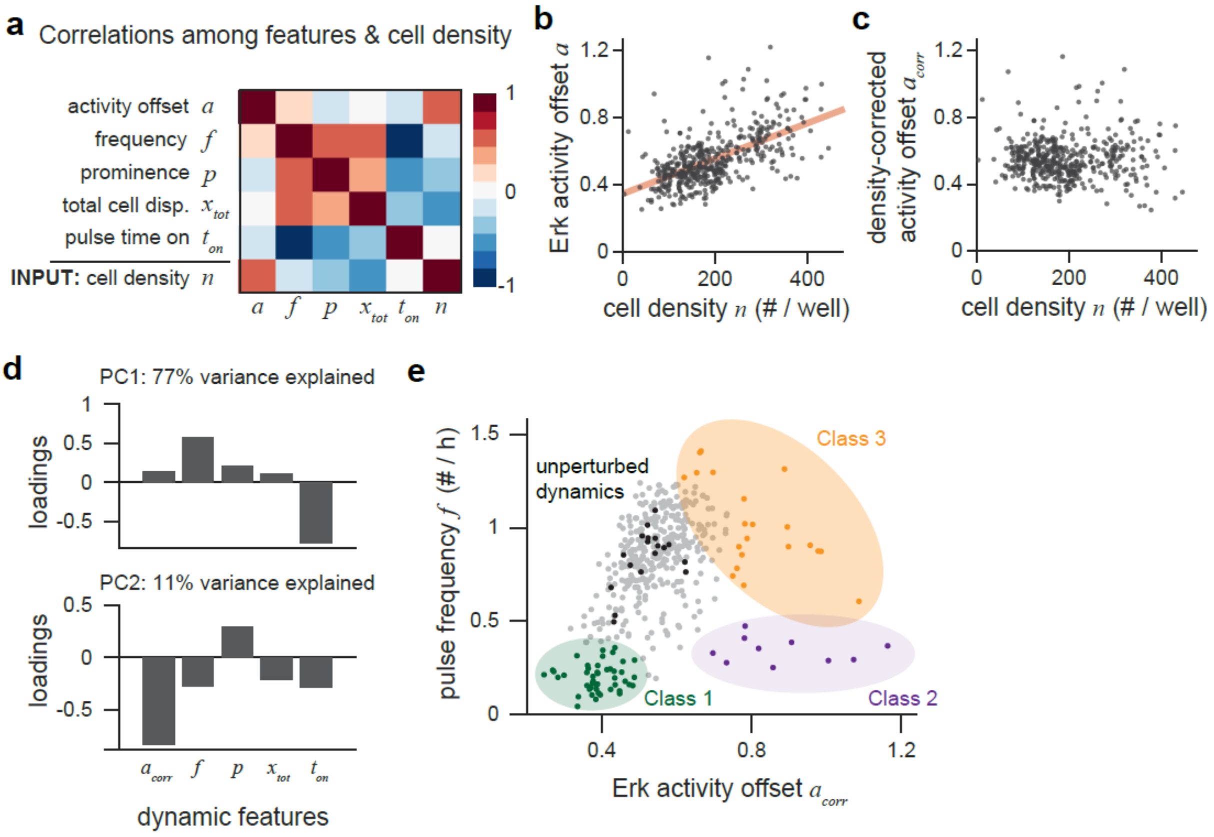

**Figure S3, related to Figure 3. Additional analyses for the signaling dynamics drug screen**. (**A**) Correlation coefficients between measurements computed across responses from all compounds in the small molecule screen. In addition to the four dynamic parameters ($a$, $f$, $p$, $t_{on}$), correlations were also measured for cells' total displacement ($x_{tot}$) as well as one independent, experimentalist-controlled variable: cell density ($n$). The high correlation observed between cell density $n$ and activity offset $a$ suggested that a correction could separate drug-induced changes in Erk activity from those dependent on cell density, a drug-independent variable. (**B**) Plot of Erk activity $a$ as a function of cell density $n$, with best-fit line shown in red. (**C**) Plot of density-corrected Erk activity offset $a_{corr}$ as a function of cell density $n$, computed by subtracting the expected offset value based on cell density (red line in **B**) from the measured offset and adding a constant value to ensure that $\langle a_{corr} \rangle = \langle a \rangle$. (**D**) Plots of the contribution of each dynamic feature onto the first two principal components. PC1 is dominated by time-related parameters (frequency $f$ and time on $t_{on}$), whereas PC2 is dominated by amplitude-related parameters (Erk offset $a_{corr}$ and pulse prominence $p$). (**E**) Plot of drug screen hits from all three classes on a plot of Erk activity offset $a_{corr}$ versus pulse frequency $f$, demonstrating that all three classes of response can be separated by these two parameters.

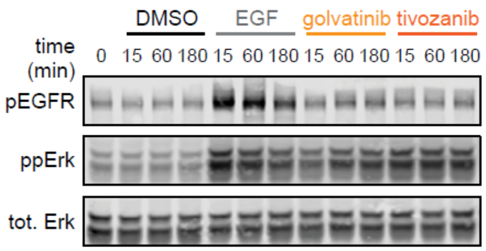

**Figure S4, related to Figure 4. Gel images for assessing drug-induced changes to Erk and EGFR phosphorylation.** A single gel was split and incubated with rabbit anti-phospho-EGFR (pEGFR), rabbit anti-dually-phosphorylated Erk (ppErk), and mouse anti-total Erk as indicated. Lysates were collected at indicated timepoints after stimulation with EGF, golvatinib, tivozanib, or DMSO as a loading control. Quantification of the Western blot is presented in **Figure 4D-E**.

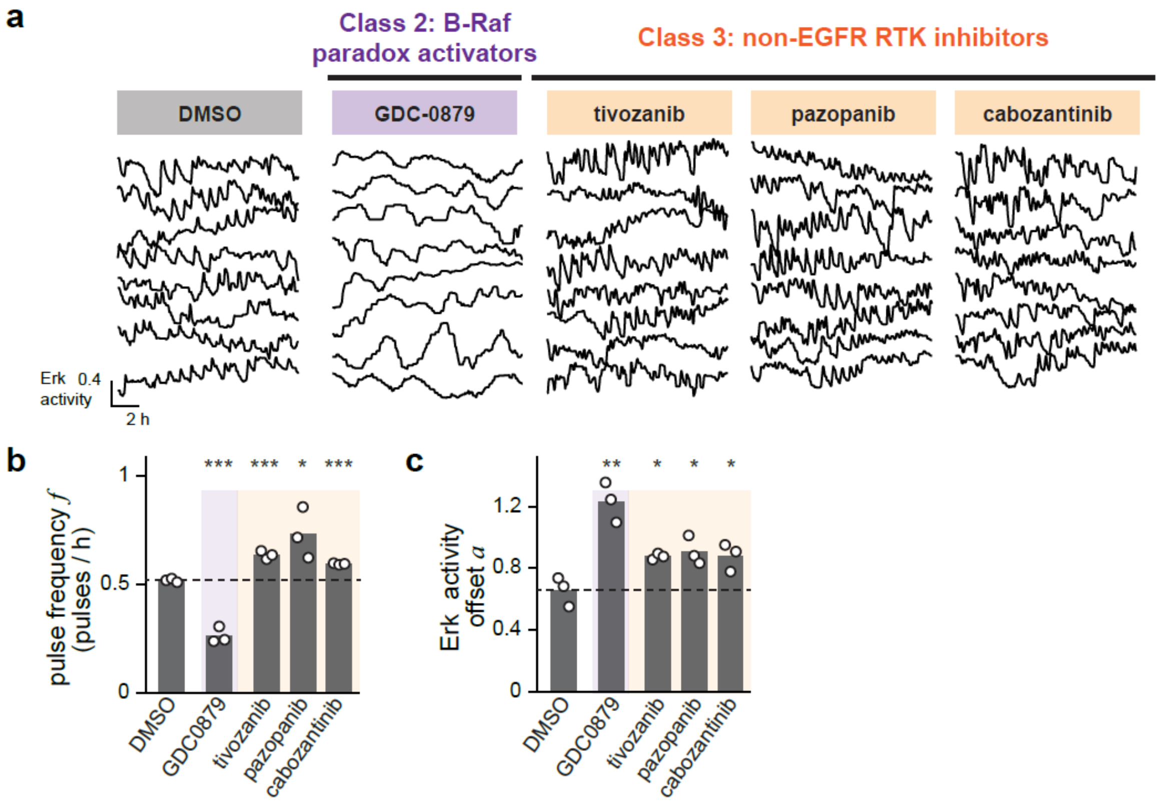

**Figure S5, related to Figure 5. Drug-altered Erk dynamics from keratinocytes cultured in complete media.** (**A**) Representative traces from keratinocytes grown in complete media that were treated with either a Class 2 B-Raf inhibitor, a Class 3 RTK inhibitor, or DMSO as a control. Cells were imaged every 3 min over 12 h. (**B-C**) Average Erk pulse frequencies (in **B**) and overall Erk activity offset (in **C**) for keratinocytes treated as in **A**. Each biological replicate (white circle) represents dynamics assessed from at least 100 single cells. Statistics are derived using a two-sided t-test (*, $p < 0.05$; **, $p < 0.01$; ***, $p < 0.001$).

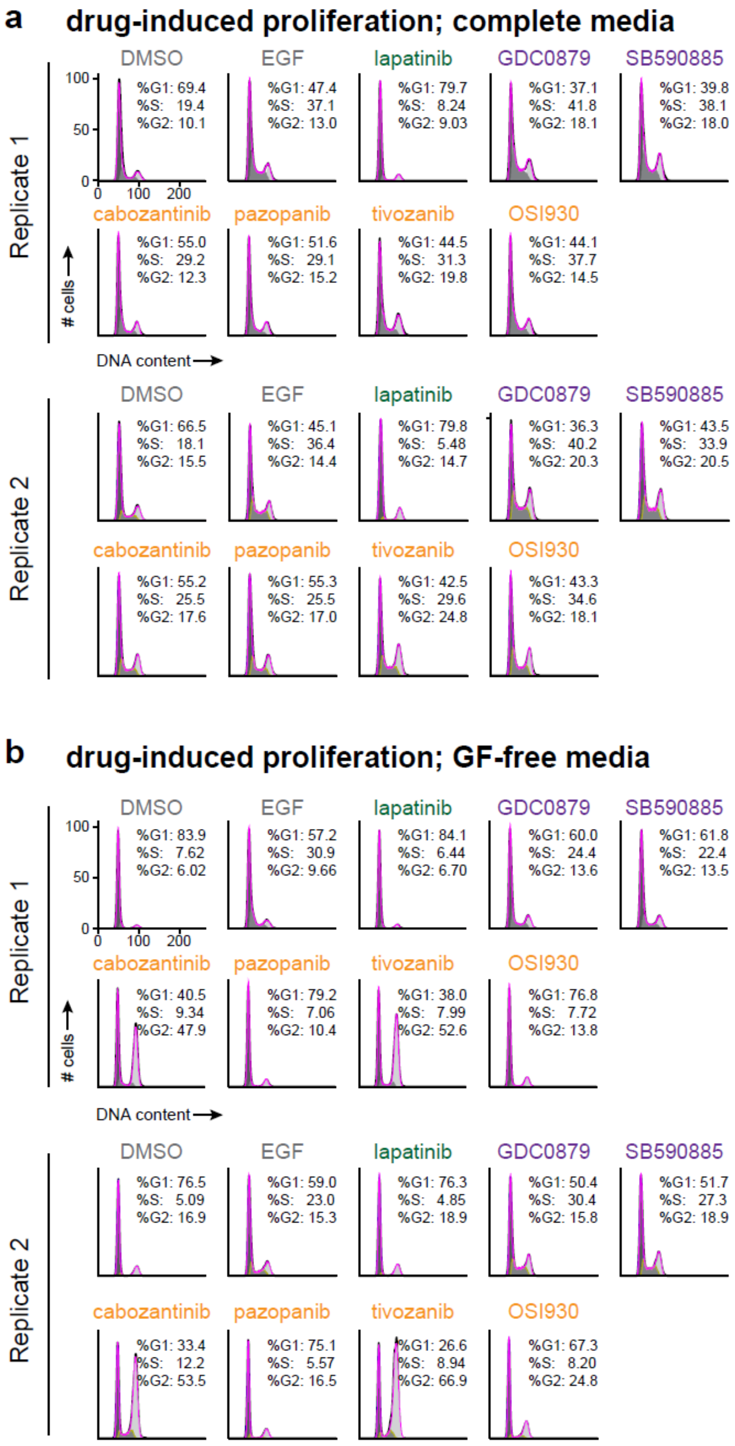

**Figure S6, related to Figure 5. Cell proliferation in response to Erk dynamics-altering drugs**. (**A**) DNA content distributions from keratinocytes grown in complete media for 22 h in the presence of selected Class 1, 2, or 3 kinase inhibitors. Two biological replicates are shown, with each stimulus condition containing data from at least 20,000 single cells. (**B**) DNA content distributions from keratinocytes grown in GF-free media for 22 h in the presence of selected Class 1, 2, or 3 kinase inhibitors. Two biological replicates are shown, with each stimulus condition containing data from at least 20,000 single cells.

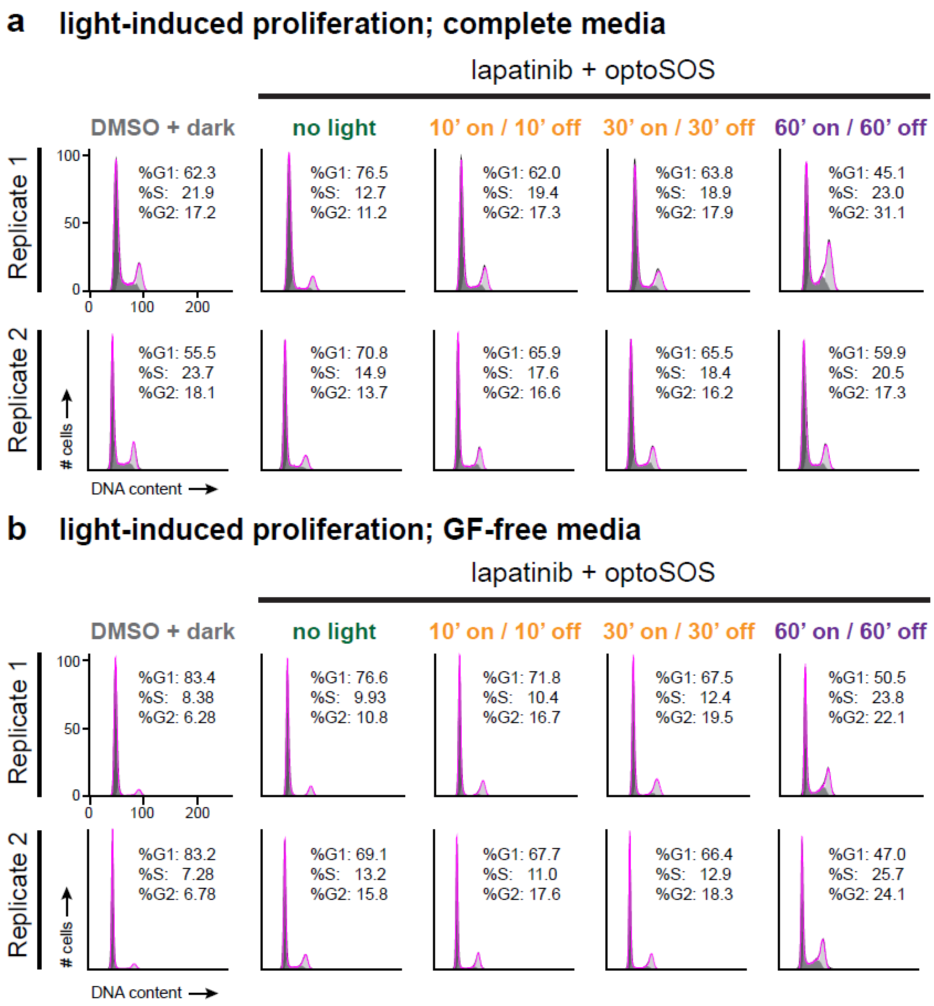

**Figure S7, related to Figure 6. Cell proliferation in response to dynamic optogenetic Ras/Erk stimuli**. (**A**) DNA content distributions from OptoSOS keratinocytes grown in complete media for 22 h and stimulated with light inputs mimicking the Erk dynamics induced by Class 1, 2, or 3 kinase inhibitors. Two biological replicates are shown, with each stimulus condition containing data from at least 20,000 single cells. (**B**) DNA content distributions from OptoSOS keratinocytes grown in GF-free media for 22 h and stimulated with light inputs mimicking the Erk dynamics induced by Class 1, 2, or 3 kinase inhibitors. Two biological replicates are shown, with each stimulus condition containing data from at least 20,000 single cells.

#### Supplementary Tables

**Supplementary Table 1: Class 1, 2, and 3 drugs and their validated targets.** Targets were obtained from the LINCS database (Keenan et al., 2018; Moret et al., 2019) and from Selleck Chemicals' annotation of their kinase inhibitor compound library. Similar isoforms were condensed into a single target class (e.g., MEK1 and MEK2: MEK; AurKA, AurKB, AurKC: AurK). From the Validated Targets list, targets within each class that function as EGFR/MEK/ERK inhibitors were annotated green, paradox-activating B-Raf inhibitors were annotated purple, and non-ErbB RTK inhibitors were annotated yellow.

| | Product Name | Validated Targets | EGFR / MEK / ERK | RAF | Non-ErbB RTK |
|---|---|---|---|---|---|
| CLASS 1 | Selumetinib (AZD6244) | MEK | | | |
| | Afatinib (BIBW2992) | ABL, BLK, DYRK EPHA, ERBB, GAK, HIPK, IRAK, LCK | | | |
| | Dasatinib | ABL, BLK, BTK, CSF1R, DDR, EPHA, EPHB, ERBB, FGR, FRK, FYN, GAK, JAK, KIT, LCK, LYN, MAP3K19, MAP4K5, MINK1, MKK5, PDGFR, RAF, RET, RIPK, SLK, SRC, TKT, TNK, TYK2, YES1 | | | |
| | Gefitinib (ZD1839) | ABL, CDK, CSNK, EPHA, ERBB, GAK, HIPK, IRAK, LCK, LYN, MAP3K19, MKK5, MKNK, RIPK, SLK, STK10 | | | |
| | Lapatinib (GW-572016) Ditosylate | ERBB, RAF | | | |
| | PD0325901 | MEK | | | |
| | AC480 (BMS-599626) | ERBB | | | |
| | SL-327 | MEK | | | |
| | Refametinib (RDEA119, Bay 86-9766) | MEK | | | |
| | WZ3146 | ERBB | | | |
| | WZ8040 | ERBB | | | |
| | CUDC-101 | ERBB | | | |
| | Pelitinib (EKB-569) | ABL, AXL, BLK, CSNK, ERBB, FGR, FRK, GAK, JAK, LCK, LYN, MAP4K5, MEK, SLK, SRC, STK10, YES1 | | | |
| | Pimasertib (AS-703026) | MEK | | | |

| | |
|---|---|
| AEE788 (NVP-AEE788) | ERBB, VEGFR |
| AZD7762 | ABL, ALK, AXL, BLK, CSF1R, DDR, DYRK, EPHA, ERBB, FLT3, FYN, IRAK, JAK, KIT, LCK, MAP3K19, MEK, MKK5, MKNK, NUAK, PDGFR, PKC, PKN, PLK, PTK2, RET, SLK, SRC, STK10, TRK, VEGFR |
| PD318088 | MEK |
| Sapitinib (AZD8931) | ERBB |
| OSI-420 | ERBB |
| TAK-733 | MEK |
| Trametinib (GSK1120212) | MEK |
| Ibrutinib (PCI-32765) | AURK, BLK, BTK, EPHB, ERBB, FGFR, JAK, LCK, MKK5, NUAK, PKD, RET, RIPK, SRC, TIE, TRK, VEGFR, YES1 |
| Tyrphostin AG-1478 | ABL, ERBB, GAK, MKNK |
| TAK-285 | ERBB |
| WHI-P154 | ERBB, JAK |
| Icotinib | ERBB |
| Binimetinib (MEK162, ARRY-162, ARRY-438162) | MEK |
| Osimertinib (AZD9291) | ERBB |
| Ulixertinib (BVD-523, VRT752271) | ERK |
| AZD3759 | ERBB |
| Cobimetinib (GDC-0973, RG7420) | MEK |
| Olmutinib (HM61713, BI 1482694) | ERBB |
| Saracatinib (AZD0530) | ABL, KIT, LCK, SRC, YES1 |
| SP600125 | JNK, MKNK |
| Hesperadin | AURK |
| TWS119 | GSK3 |

| | | |
|---|---|---|
| | AZ 960 | JAK |
| | Volasertib (BI 6727) | PLK |
| | Omipalisib (GSK2126458, GSK458) | MTOR, PIK3C |
| | WYE-125132 (WYE-132) | MTOR |
| | NVP-BSK805 2HCl | JAK |
| | ZM 306416 | VEGFR |
| | Ro3280 | PLK |
| CLASS 2 | GDC-0879 | CSNK, MAP3K19, RAF |
| | SB590885 | RAF |
| | Dovitinib (TKI-258, CHIR-258) | ABL, AMPK, AURK, BLK, CDK, CSF1R, CSNK, EPHB, FGFR, FGR, FLT3, FYN, GAK, GSK3, HIPK, IKBK, IRAK, JAK, KIT, LCK, LYN, MAP3K19, MAP4K5, MEK, MINK1, MKK5, MYLK, NUAK, PDGFR, PKN, PLK, RET, RIPK, ROCK, RPS6KA, SLK, STK10, TRK, VEGFR, YES1 |
| | Dovitinib (TKI-258) Dilactic Acid | ABL, AMPK, AURK, BLK, CDK, CSF1R, CSNK, EPHB, FGFR, FGR, FLT3, FYN, GAK, GSK3, HIPK, IKBK, IRAK, JAK, KIT, LCK, LYN, MAP3K19, MAP4K5, MEK, MINK1, MKK5, MYLK, NUAK, PDGFR, PKN, PLK, RET, RIPK, ROCK, RPS6KA, SLK, STK10, TRK, VEGFR, YES1 |
| | Rebastinib (DCC-2036) | ABL, FGR, FLT3, LYN, SRC, TIE, VEGFR, YES1 |
| | A-674563 | AKT, CDK |
| | AZD8055 | MTOR |
| | LY2090314 | AMPK, CDK, CLK, GSK3, LCK, PKC, PKN |
| | abemaciclib (LY2835219) | CDK |
| | SC1 | ERK |
| | SGI-7079 | VEGFR |
| | Pazopanib HCl (GW786034 HCl) | ABL, AURK, CSF1R, DDR, EPHB, FGFR, FRK, IRAK, KIT, LYN, MAP3K19, MKK5, PDGFR, PLK, RAF, RET, RIPK, SLK, STK10, TIE, VEGFR |

| | | |
|---|---|---|
| CLASS 3 | ZM 447439 | AURK, ERBB, FYN, LCK, MEK, PDGFR, RET, ROCK, VEGFR |
| | Danusertib (PHA-739358) | ABL, AURK, FGFR, RET |
| | Foretinib (GSK1363089) | ABL, AMPK, AURK, BLK, CDK, CLK, CSF1R, DDR, EPHA, EPHB, ERBB, FGFR, FLT3, FRK, FYN, HIPK, IRAK, JNK, KIT, LCK, LYN, MAP3K19, MEK, MET, MKK5, PDGFR, PLK, RET, RIPK, SLK, SRC, STK10, TIE, TRK, VEGFR, YES1 |
| | JNJ-38877605 | MET |
| | Cabozantinib (XL184, BMS-907351) | FLT3, KIT, MET, RET, TIE, TRK, VEGFR |
| | BMS-754807 | MET, TRK |
| | AT9283 | ABL, AURK, FGFR, FLT3, GSK3, JAK, RET, S6K, VEGFR, YES1 |
| | Barasertib (AZD1152-HQPA) | AURK, EPHB, ERBB, FLT3, HIPK, KIT, LCK, MKK5, PDGFR, RET, TIE, VEGFR |
| | PLX-4720 | CDK, EPHB, MEK, MKK5, PDGFR, RAF, RIPK |
| | CYC116 | AURK, VEGFR |
| | Regorafenib (BAY 73-4506) | FGFR, KIT, PDGFR, RAF, RET, TIE, VEGFR |
| | CHIR-99021 (CT99021) | ABL, FYN, GSK3, VEGFR |
| | PD173074 | FGFR, VEGFR |
| | Golvatinib (E7050) | MET, VEGFR |
| | 7,8-Dihydroxyflavone | TRK |
| | LY294002 | CSNK, GSK3, MTOR, PIK3C |
| | OSU-03012 (AR-12) | PDK |
| | SNS-032 (BMS-387032) | CDK, CLK, CSNK, GSK3 |
| | PD98059 | MEK |
| | AT7867 | S6K |
| | Uprosertib (GSK2141795) | AKT |
| | AT13148 | ROCK, S6K |
| | Butein | ERBB |
| | ETP-46464 | MTOR |

#### Supplementary Video Legends

**Movie S1.** Time-lapse imaging of primary mouse keratinocytes expressing the H2B-RFP nuclear marker (not shown) and KTR-BFP Erk activity biosensor (shown). Cells were imaged using a 20X air objective every 2 min for 24 h under continuous culture in GF-free media, and exhibit repeated Erk pulses throughout the video. Timer indicates hh:mm; scale bar indicates 30 µm. Related to Figure 2.

**Movie S2.** Time-lapse imaging of MCF10A human breast epithelial cells and primary mouse keratinocytes expressing the H2B-RFP nuclear marker (not shown) and KTR-BFP Erk activity biosensor (shown). Cells were imaged using a 10X air objective every 3 min for 4 h under continuous culture in GF-free media supplemented with different doses of EGF (indicated above). Timer indicates hh:mm; scale bar indicates 30 µm. Related to Figure 2.

**Movie S3.** Time-lapse imaging of primary mouse keratinocytes expressing the H2B-RFP nuclear marker (not shown) and KTR-BFP Erk activity biosensor (shown). Cells were imaged using a 10X air objective every 3 min for 5 h under continuous culture in GF-free media supplemented with the kinase inhibitors indicated at a concentration of 2.5 µM. Timer indicates hh:mm; scale bar indicates 30 µm. Related to Figure 3.

**Movie S4.** Time-lapse imaging of primary mouse keratinocytes expressing the H2B-RFP nuclear marker (not shown) and KTR-BFP Erk activity biosensor (shown). Cells were imaged using a 20X air objective every 3 min for 16 h under continuous culture in GF-free media supplemented the kinase inhibitors indicated at a concentration of 2.5 µM or neutralizing antibodies against Met or VEGFR2. Timer indicates hh:mm; scale bar indicates 30 µm. Related to Figure 4.

**Movie S5.** Time-lapse imaging of primary mouse keratinocytes expressing the OptoSOS system (not shown), H2B-RFP nuclear marker (not shown) and KTR-iRFP Erk activity biosensor (shown). Cells were imaged using a 20X air objective every 90 sec for 15 h under continuous culture in GF-free media. At t = 2 h (as indicated by the "+EGFRi" label) cells were treated with 2.5 µM lapatinib. At t = 3 h, cells were stimulated with 15 min pulses of 450 nm blue light. Blue box indicates times of light delivery. Timer indicates hh:mm; scale bar indicates 30 µm. Related to Figure 6.
